# Correlated STORM-homoFRET Imaging: Applications to the Study of Membrane Receptor Self-Association and Clustering

**DOI:** 10.1101/2021.12.21.473581

**Authors:** Amine Driouchi, Scott D. Gray-Owen, Christopher M. Yip

**Affiliations:** Department of Biochemistry, University of Toronto, 1 King’s College Circle, Medical Sciences Building, Toronto, ON, M5S1A8, Canada; Department of Molecular Genetics, University of Toronto, 1 King’s College Circle, Medical Sciences Building, Toronto, ON, M5S1A8, Canada; Institute of Biomedical Engineering, Terrence Donnelly Centre for Cellular and Biomolecular Research, University of Toronto, Toronto, ON, M5S 3E1, Canada; Department of Chemical Engineering & Applied Chemistry, 200 College St, Toronto, ON, M5S 3E5, Canada; Terrence Donnelly Centre for Cellular & Biomolecular Research, University of Toronto, 160 College Street, Toronto, Ontario M5S 3E1, Canada

**Keywords:** Super-resolution microscopy, STORM, homoFRET, membrane proteins, oligomers, nanobodies, clustering

## Abstract

Mapping the self-organization and spatial distribution of membrane proteins is key to understanding their function. We report here on a correlated STORM/homoFRET imaging approach for resolving the nanoscale distribution and oligomeric state of membrane proteins. Live cell homoFRET imaging of CEACAM1, a cell-surface receptor known to exist in a complex equilibrium between monomer and dimer/oligomer states, revealed highly heterogenous diffraction-limited structures on the surface of HeLa cells. Correlated super-resolved STORM imaging revealed that these structures comprised a complex mixture and spatial distribution of self-associated CEACAM1 molecules. This correlated approach provides a compelling strategy for addressing challenging questions about the interplay between membrane protein concentration, distribution, interaction, clustering, and function.

## Introduction

Identifying the factors that govern the spatial-temporal distribution and interaction of membrane proteins is critical for addressing key questions in cell biology. These include determining the structural underpinnings of intercellular engagement and the mechanisms of effector signaling upon receptor activation. Identifying the key structure-function relationships relies upon quantifying protein dynamics and association state [1–4]. Techniques such as Förster resonance energy transfer (FRET) [5–7] and fluorescence correlation spectroscopy (FCS) [8, 9] are often employed to quantify and spatially map protein-protein interactions, clustering, self-association, and dynamics. One commonly used approach for characterizing self-association is homoFRET, which occurs due to energy transfer between identical fluorophores that can act as either donors or acceptors. In homoFRET, non-radiative energy transfer between identical fluorophores located within the requisite Förster distance results in depolarized emission upon excitation with linearly polarized light. As such, this method reports on interactions that occur over distances less than 10 nm and with transfer efficiencies dependent on the fluorophore, local structure, and extent of conformational flexibility [5, 6, 10]. The result of homoFRET imaging is a diffraction-limited pixel-wise anisotropy map that reports the spatial distribution of self-association.

Although FRET-based approaches provide insights into the proximity of two interacting fluorophores, the diffraction limit classically limits the extent of spatial localization. Recent developments in super-resolution fluorescence microscopy or single molecule localization microscopy (SMLM) are allowing researchers to resolve objects with a localization precision of 20 to 100 nm, often revealing sub-diffraction limit structural features [11, 12]. These techniques can be subdivided into two categories: ensemble imaging approaches, which include stimulated emission depletion (STED)[1, 13–16] and structured illumination (SIM) microscopies [16–19], and single molecule localization microscopy (SMLM), which includes stochastic optical reconstruction microscopy (STORM) [20–22] and photoactivated localization microscopy (PALM) [23]. A key limitation for SMLM lies in determining whether the localized structures are in fact interacting or are simply clustered in a non-interacting fashion. This is further complicated by the very nature of the SMLM acquisition and image processing algorithms, which typically rely on stochastic excitation of, and emission from, individual fluorophores. This challenge prompted us to develop an integrated approach that would marry the high spatial resolution of SMLM with self-association data provided by homoFRET. Since homoFRET images are diffraction-limited, the pixel-wise *r*-values can only report on the average association state within the pixel, and not the true spatial distribution. In principle, correlating super-resolution and homoFRET images would provide new insights into the relationships between oligomeric state and spatial distribution [24]. Here we describe the methodology behind a correlative STORM/homoFRET technique (**Figures 1, S1**). When applied to membrane or membrane-associated proteins, this correlated approach can reveal their nano-scale distribution and oligomerization or self-association state.

**Figure 1.**
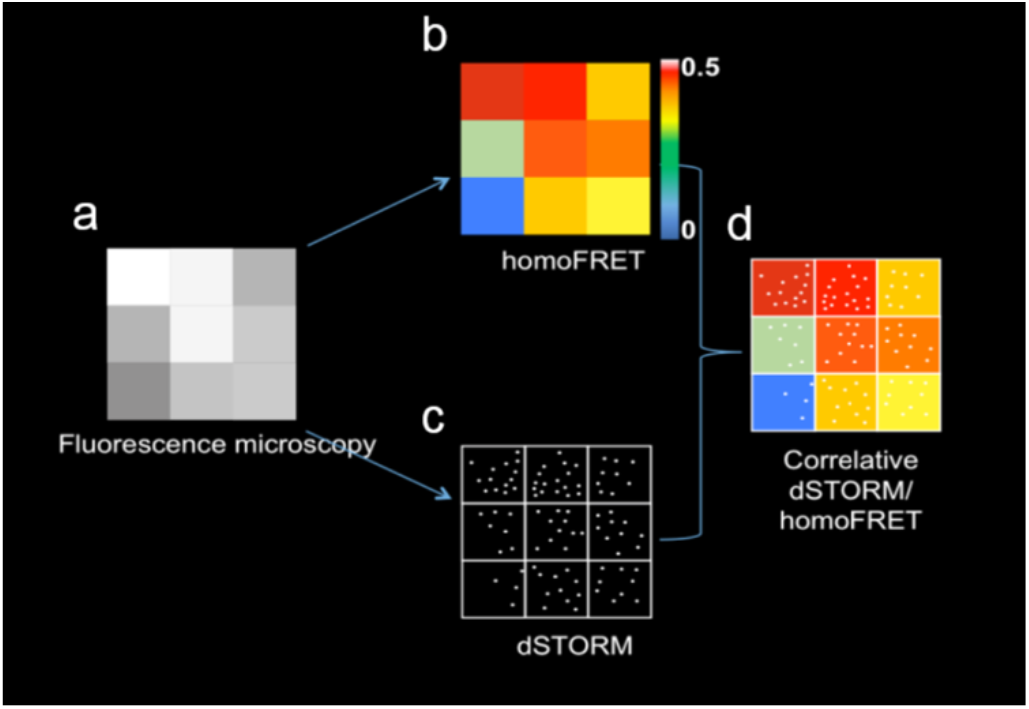
Schematic representation of the correlated STORM/homoFRET approach using a model 3 x 3 pixel image. **(a)** Diffraction-limited fluorescence microscopy with each pixel displaying a grey-scale intensity value. **(b)** In a homoFRET image, each pixel is assigned an *r*-value representing the pixel’s average anisotropy, with lower values reflecting increased selfassociation. **(c)** An idealized STORM image showing individual localizations without their respective localization uncertainty. **(d)** Correlated STORM/homoFRET allows matching of the true spatial distribution with the pixel-wise anisotropy values.

### CEACAM1-4L, a transmembrane cell-adhesion molecule

For the purposes of the present work, we focused on the family of carcinoembryonic antigen-related cellular adhesion molecules (CEACAMs), which are cell surface glycoproteins involved in homo- and hetero-philic intercellular interactions involved in cellular growth, differentiation, tumourigenesis, inflammation, and infection [25]. Despite CEACAMs’ importance in health and disease, the nature of CEACAM interactions and their mechanisms of signaling remain poorly understood. The full-length splice variant of CEACAM1, the evolutionary progenitor of the CEACAM family, has an extracellular domain consisting of an amino-terminal IgV-like domain and three IgC2-like domains, with all these domains being heavily glycosylated [26, 27]. This protein is anchored into the membrane via a transmembrane sequence and a 76-amino acid cytoplasmic domain that contains two immunoreceptor tyrosine-based inhibitory motifs (ITIMs), although other variants lack these sequences [25]. While it has been known that CEACAM1 can exist as a monomer, a dimer, or higher order oligomer, and that different oligomeric forms engage different downstream signaling cascades, how these complexes are distributed remains unknown [28–30] **(Figure 2)**. Furthermore, the dynamics and interchange of these oligomeric states remain poorly characterized. These questions prompted us to develop a correlative STORM/homoFRET-based approach for characterizing the nanoscale distribution of CEACAM1-4L monomers and oligomers on the cell surface. In this study, homoFRET measurements were performed using YFP-labelled CEACAM1-4L while STORM imaging was accomplished through a camelid anti-YFP single domain antibody labeled with Alexa Fluor 647 (AF647).

**Figure 2.**
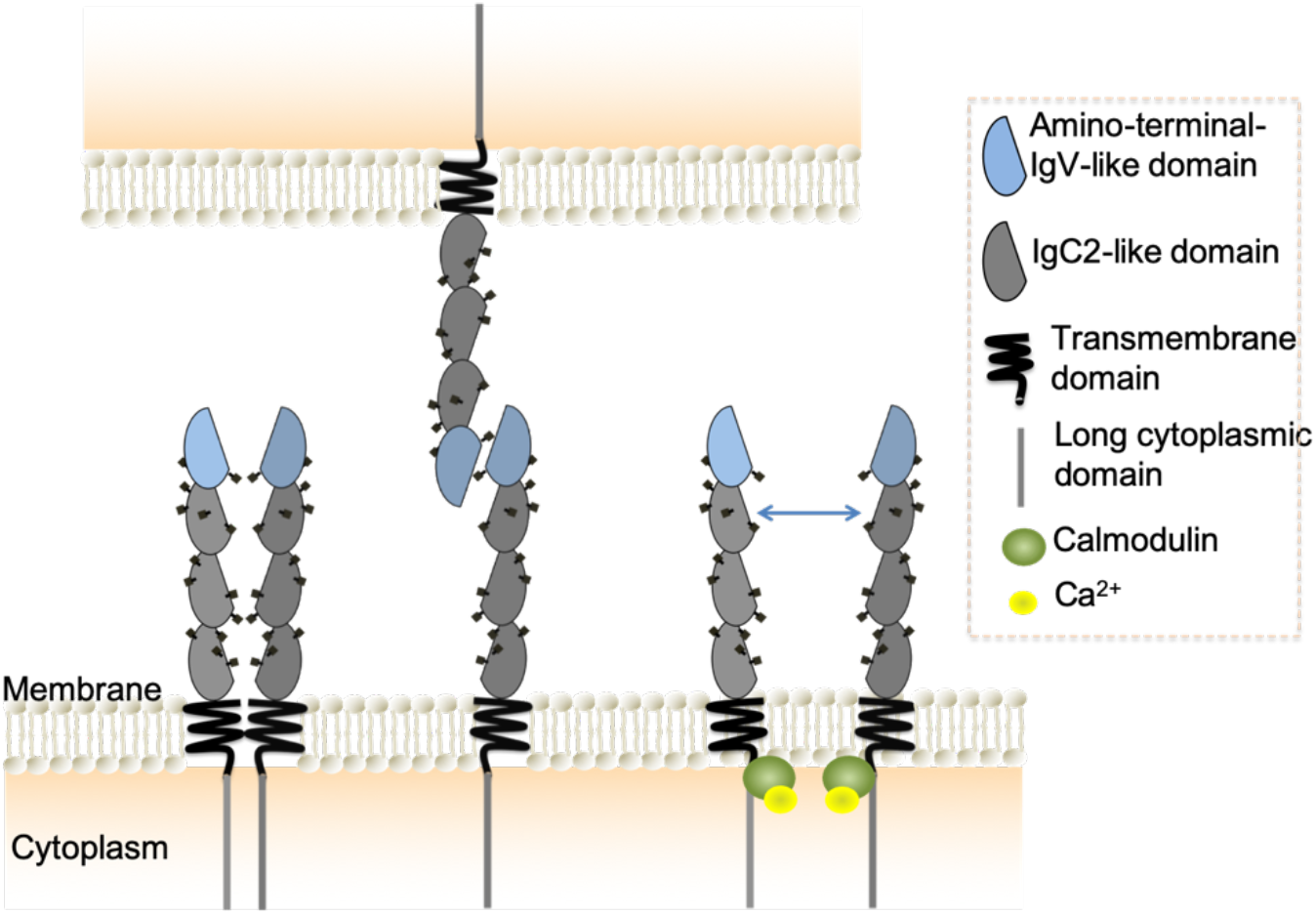
CEACAM1-4L is known to exist as a monomer, dimer, or higher-order oligomer. This equilibrium is regulated by key cofactors, including disruption by binding of calcium-loaded calmodulin to the cytoplasmic domain.

## Results

### Steady-state fluorescence anisotropy of CEACAM1-eYFP

The human endocervically-derived HeLa cells are one of the few epithelial cell lines that lacks endogenous CEACAMs, allowing ectopic expression of individual CEACAM1-4L family members to ascribe function. Past work has shown that ectopically expressed CEACAM1-4L (CEACAM1) can transition between the monomeric and cis-dimeric forms seen in cell lines that naturally express this protein [28, 31]. Thus, for these studies, we have used HeLa cells transiently transfected with a chimeric allele encoding CEACAM1 with a carboxyl-terminus fused eYFP (CEACAM1-eYFP).

To establish the range of values attainable, TIRF-based homoFRET measurements were determined using HeLa cells expressing soluble monomeric Venus and the tandemly-expressed dimeric Venus (**Figure S4).** Before this and all subsequent experiments, the G-factor correction factor was measured to correct for potentially varying light sensitivities of the detectors capturing parallel and perpendicular polarizations. In addition, we have shown that the resulting pixel-wise anisotropy map is independent of the given intensity values and is therefore a ratio between the two captured emission channels **(Figure S5)**.

Live cell TIRF-homoFRET imaging of CEACAM1-eYFP in HeLa cells revealed the presence of both diffuse and irregularly bright regions. Counterintuitively, the low intensity/diffuse regions were comprised of predominantly dimeric or oligomeric CEACAM1 with anisotropy values ranging from 0.1 to 0.2, while the high intensity/clustered regions appeared more monomeric with anisotropy values ranging from 0.3 to 0.45 **(Figure 3a).** Saturated pixels are not taken into consideration in the homoFRET calculation as the anisotropy calculation requires two intensity values from the collected images while low intensity pixels are removed using a threshold value kept constant for all analyzed cells **(Figure 3d).** The presence of cell-to-cell variability suggests that the observed equilibrium is regulated by the cell (**Figure S2).** In addition, we performed homoFRET experiments on a well characterized CEACAM1 mutant known to be unable to form cis-dimers. These experiments revealed a dramatic shift to higher anisotropy values for this monomeric mutant (**Figure S6**). We subsequently performed homoFRET experiments on a CEACAM1 mutant known to be unable to form trans-homophilic interactions. In this case, the histogram distribution is shifted towards lower pixel values indicative of enrichment in cis-dimers and oligomers **(Figure S6).** Although not the focus of this paper, these insights highlight the fact that most oligomeric CEACAM1 exists likely in the cis conformation. Overall, these experiments highlight the utility of this approach for studying self-association states.

**Figure 3.**
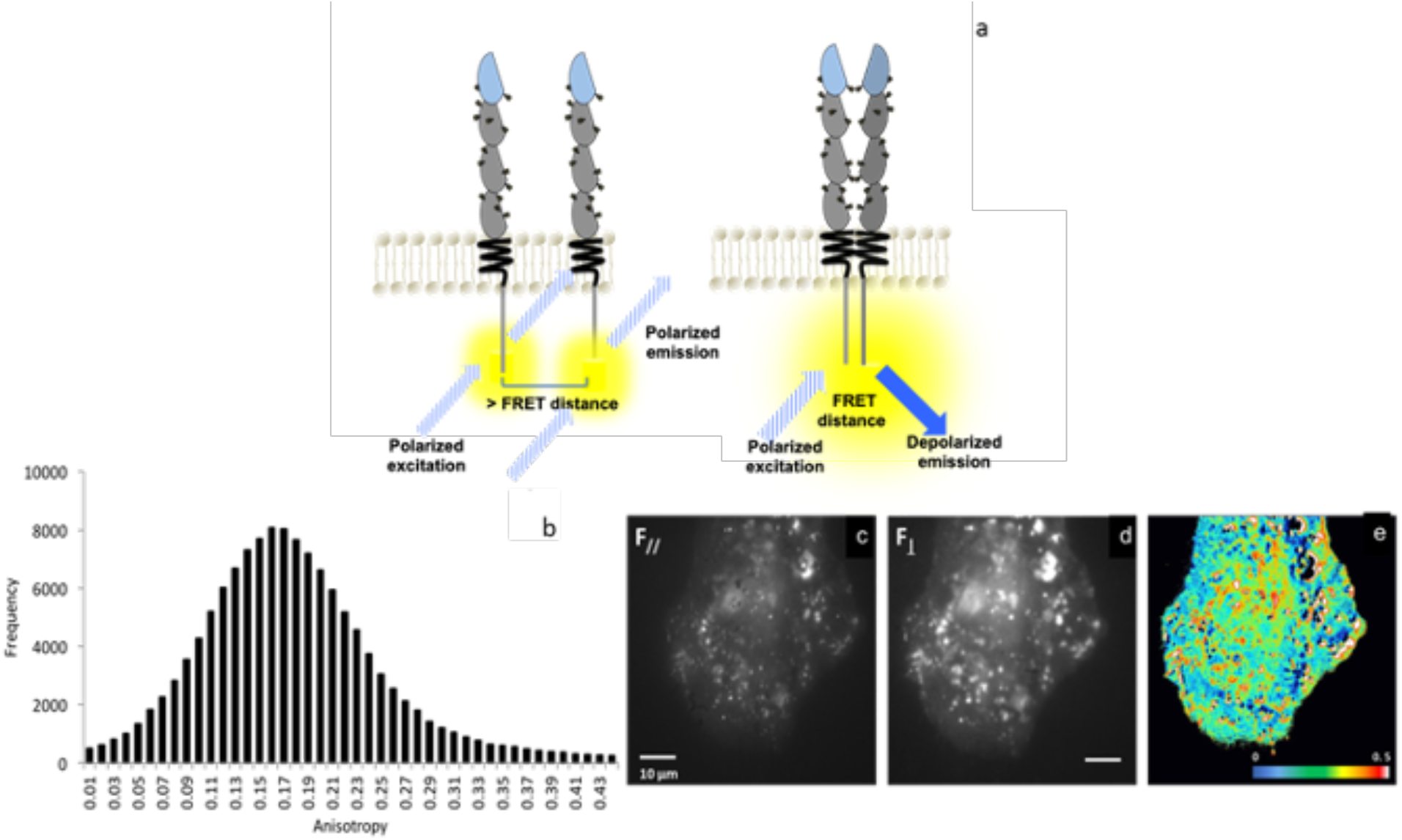
HomoFRET provides an anisotropy map representative of the distribution of monomers and oligomers. **(a)** Schematic showing depolarized emission as a result of FRET when two identical fluorophores are within the requisite FRET distance. **(b)** Distribution of anisotropies within HeLa cell transiently transfected with CEACAM1-eYFP. **(c)** Polarized (F_ll_) diffraction-limited image. **(d)** Depolarized (F_⊥_) diffraction-limited image. **(e)** Anisotropy map illustrating the heterogeneous distribution of CEACAM1 monomers and oligomers, corresponding to high (red) and low anisotropy values (blue), respectively.

There is clearly a non-uniform spatial distribution of anisotropies across the surface of each cell, with an inverse correlation between CEACAM1 density (reflected by eYFP intensity) and the proportion of CEACAM1 that is in an oligomeric state (reflected by lower anisotropy values). However, the diffraction-limited spatial resolution of homo-FRET prevents clear characterization of CEACAM1 organization within these regions. To overcome this limitation, we applied STORM imaging to the same samples that had been characterized by TIRF-homoFRET imaging.

### Single domain antibody labeling for STORM

For STORM imaging, CEACAM1-eYFP is labeled with an AF647 labelled anti-eYFP camelid single domain antibody **(Figure 4c)** [32][33]. A key challenge in the use of antibodies for STORM imaging lies in the size of the antibodies themselves, which are typically ~ 13 nm long. This can result in significant variability in the localization. To address this, we elected to use a nanobody comprised solely of the 12-15 kDa variable domain of a camelid antibody heavy chain. This nanobody is ~ 1.5-2.5 nm in size and has been specifically engineered to bind to the GFP β-barrel and its derivatives (including eYFP) with high specificity and affinity [34]. Unlike secondary antibodies that are usually conjugated with 5-8 fluorophores, nanobodies only carry 1-2 fluorophores. This reduces the likelihood of over-localizing the individual antibody (or nanobody) during STORM imaging while minimizing the generation of self-clustering artifacts. Overlapping point spread functions (PSFs) can lead to mis-localization, and under- or over-estimation of single emitters. This can lead to the artifactual creation of clusters or nanodomains, even in the case of a homogeneous distribution. As described by Sauer and colleagues [46], in order to minimize false multiple-fluorophore localizations, the lifetime of the off-state has to be significantly longer than that of the on-state. In this way, the distances between individual blinks are large enough to minimize PSF overlaps. Here we aim to measure the average distances between individual PSFs for both the antibody and nanobody labeling strategies to confirm that we have achieved sufficient separation.

**Figure 4.**
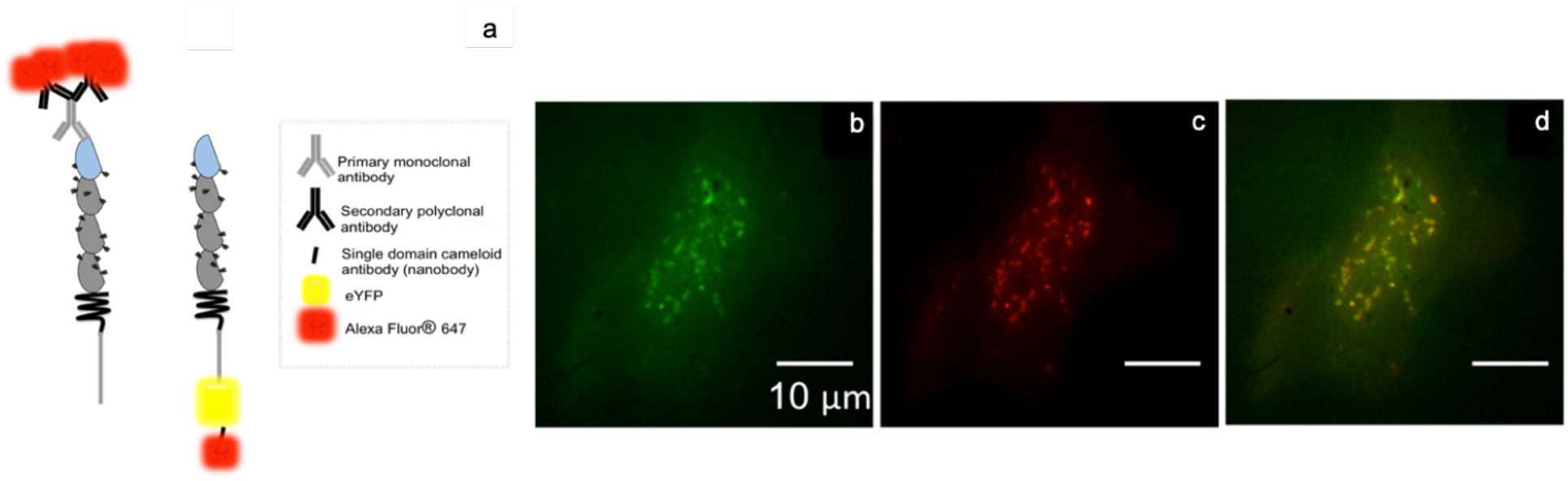
Using an eYFP-specific nanobody facilitates correlative STORM/homoFRET due to its low number of fluorophores and their close proximity to eYFP. **(a)** Schematic illustration to highlight spatial differences between CEACAM1 labeling using a monoclonal antibody targeting the extracellular domain with an AF647-labeled secondary polyclonal antibodies, versus directly conjugated single domain camelid antibody (nanobody). Each primary antibody can bind several polyclonal antibodies, and each secondary antibody typically bears 4-8 fluorophores, whereas the camelid antibodies possess 1 or 2. CEACAM1 labeling using an eYFP-specific nanobody. Representative fluorescence image of: **(b)** CEACAM1-eYFP, **(c)** CEACAM1-nanobody-AF647, **(d)** Merged eYFP and AF647 image.

We selected 10 frames from the 1000 frames dataset acquired for a given cell and super-resolved the individual frames prior to conducting a nearest-neighbour analysis on the resulting coordinates **(Figure S8).** To compare and contrast the antibody and nanobody labelling approaches, 3 replicate experiments were conducted. The average nearest neighbour distances for the antibody and nanobody labeling strategies were 1725 nm (± 19) and 2040 nm (± 26), respectively. These results suggest that both strategies generate comparable localizations and are appropriate for STORM imaging of protein nanoclusters.

For STORM imaging, the CEACAM1-eYFP HeLa cells were fixed with paraformaldehyde, permeabilized and then labeled with 1 nM of AF647-conjugated anti-YFP nanobody for 30 min **(Figure S7).** Analysis of conventional wide-field fluorescence images **(Figure 4, b-d)** revealed that the eYFP- and AF647-labeled regions were highly co-localized (Pearson coefficient = 0.861 ± 0.05 measured from 6 cells for each of 3 replicate experiments **(Figure S3).** In addition, since the AF647-labeled nanobody is bound to an eYFP which itself is bound to a flexible linker attached to the cytoplasmic domain of CEACAM1, we characterized the small, but non-negligible fluctuation of the fluorophore’s position over a 1 min acquisition period **(Figure S9).** We found this fluctuation to be well below the localization precision of STORM imaging. These data suggest that this approach provides for highly specific high-affinity labeling and thus enables correlative STORM/homoFRET experiments.

### Quantifying the impact of cell permeabilization and single domain antibody labeling on eYFP anisotropy

While homoFRET of CEACAM1-eYFP can be performed on live cells, staining with the anti-eYFP nanobody requires fixation and permeabilization of the cells. To assess how these steps may affect correlation of the anisotropy and STORM datasets, anisotropy measurements were performed after each fixation and permeabilization step **(Figure 5 ac)**. While close inspection of the corresponding images indicates that the cluster distributions and morphologies remained largely unaffected by these steps **(Figure S10)**, permeabilization did produce an apparent shift towards lower anisotropies, or oligomeric CEACAM1.

**Figure 5.**
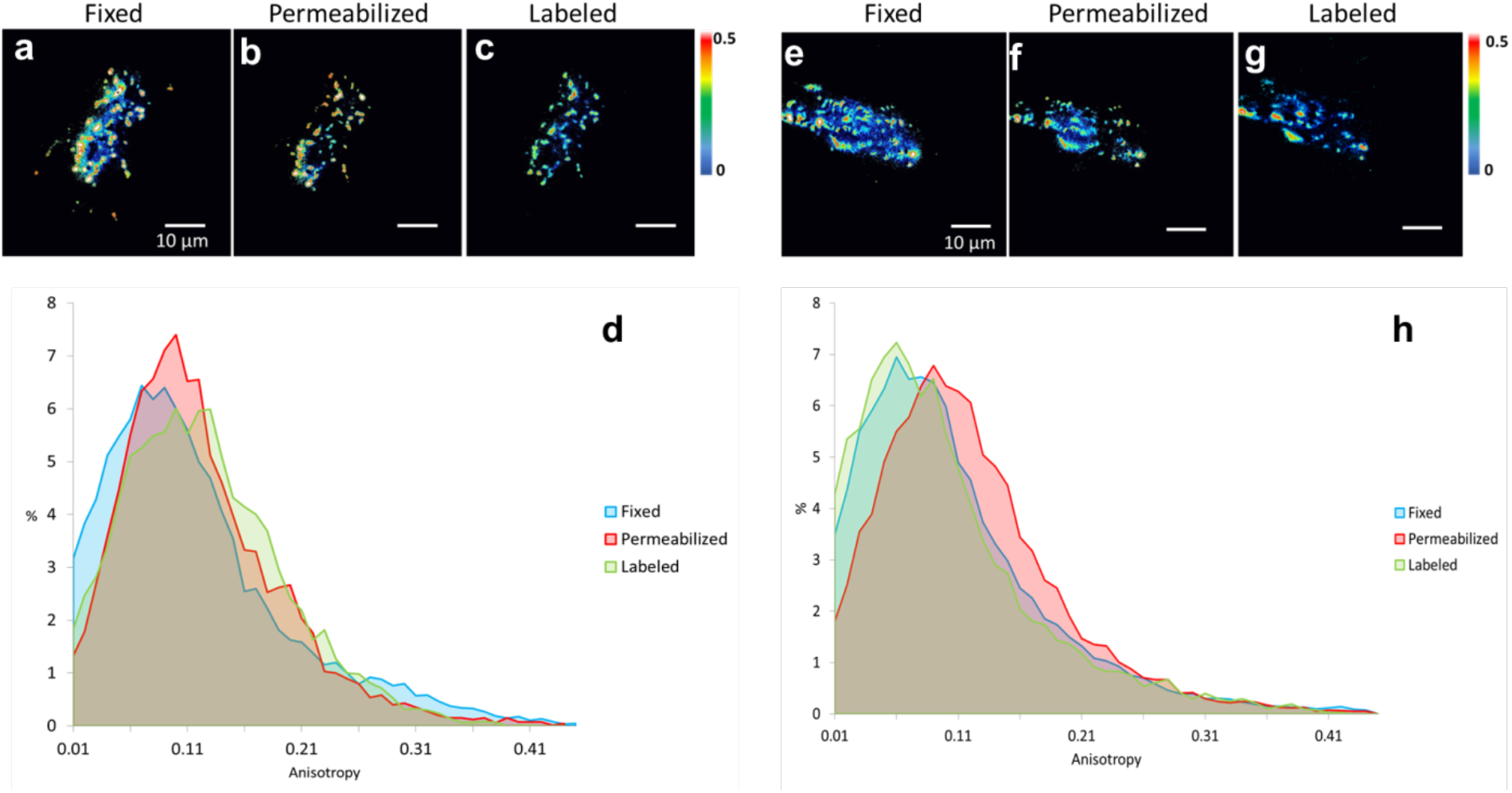
Measuring potential distortions caused by the single domain antibody-dependent immunostaining. Images depicting anisotropy measurements following cell fixation using 4% PFA **(a**, **e)**, cell permeabilization using 0.1% Triton X-100 **(b, f)**, nanobody-labeling **(c**, **g)**. Histograms **(d**, **h)** of the anisotropy distribution of the cells depicted in **a-c** and **e-g**, respectively, illustrating the overall pixel-wise change in oligomerization states upon disruption through fixation, permeabilization, and labeling.

The cumulative anisotropy plots revealed no significant shift in anisotropy upon fixation whereas permeabilization caused a small shift to the right. **(Figure 5 d, h)** However, after nanobody labeling, the cumulative anisotropy curve appeared to revert to values comparable to those originally obtained in the fixed but non-permeabilized samples. This suggests that detergent-induced distortion occurring during permeabilization may be transient in nature. Overall, fixation, permeabilization and labeling as performed in our experiments appear to minimally alter self-association states, clustering, and overall cell morphology (see Methods),

### Investigating the nanoscale organization of nanobody-labeled CEACAM1 using dSTORM

Our diffraction-limited TIRF microscopy revealed that CEACAM1 is distributed either in a diffuse manner or as clusters exhibiting various sizes and shapes. To understand the nature of the CEACAM1 species within the clusters, Voronoi tessellation-based cluster analysis was performed on super-resolved CEACAM1 datasets using the SR-Tesseller package **(Figure 6a-c).** Voronoi tessellation subdivides a set of coordinates into polygonal regions by tracing the bisector to each of two nearest neighbours. For each coordinate, the mean distance to the adjacent nearest neighbours can be calculated, generating a mean distance distribution per cell or region of interest. Regions of interest can be thresholded out and clusters identified by setting the number of localizations and cluster area minima and maxima **(Figure S11).** Remarkably, this clustering analysis revealed the presence of a consistent trimodal distribution of the mean distance histogram, suggestive of the presence of diffuse regions, microclusters, and nanoclusters **(Figure 6, S12).** Here, microclusters and nanoclusters are defined as a function of cluster density that appears to correlate well with cluster area and diameter. Similar higher density regions have been reported for other membrane proteins and are thought to exist in order to restrict their activity to intercellular contact points or to co-localize with subcellular structures [35–38].

**Figure 6.**
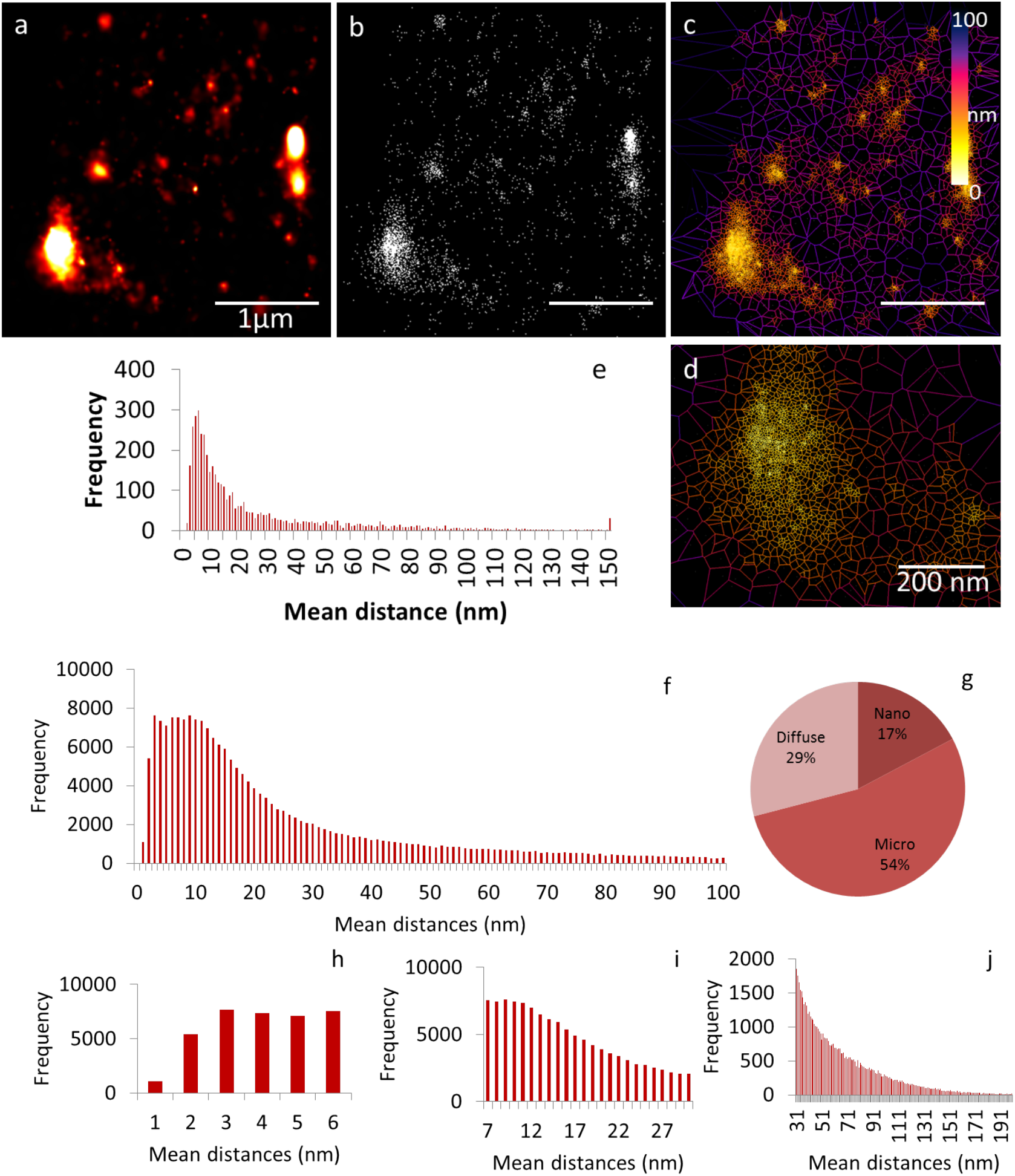
Voronoi tessellation enables distinction between three different classes of clusters. **(a)** Gaussian rendering of super-resolved CEACAM1 on HeLa cell, representative of the localization uncertainty of each point. **(b)** Scatter plot of super resolved CEACAM1. The points are presented without the localization uncertainty. **(c)** Voronoi tessellation. **(d)** Voronoi tessellation of an enlarged CEACAM1 cluster (compare scale bars in **a** and **d**). **(e)** Distribution of mean distances for tessellated ROI. **(f)** Cumulative frequency plot of mean distances collected from 4 cells. **(g)** Pie chart showing the average clustering of CEACAM1 localizations. **(h)** Nanocluster (1-6 nm) frequency plot of mean distances. **(i)** Microcluster (7-30 nm) frequency plot of mean distances. **(j)** Diffuse (>31 nm) regions frequency plot of mean distances.

Once the different cluster groups were identified, we measured area, density, circularity, and diameter for each of the nanocluster and microclustered regions from 4 cells. While circularities are similar (0.55-0.60), the average cluster diameter for microclusters and nanoclusters were 215 nm ± 27 and 49 nm ± 28, respectively **(Figure S16 b, c).** Finally, nanoclusters appear almost three times denser than microclusters (**Figure S16d**). These differences may reflect functional differences between the microclusters and nanoclusters and differences in their association to lipid membrane and cytoskeletal components.

### Correlating homoFRET and dSTORM

After cluster analysis via Voronoi tessellation, the diffraction-limited homoFRET image was scaled up without interpolation to a 6502 x 6502 pixel 16-bit image and then overlaid on the corresponding STORM image. CEACAM1 clusters of different shapes, sizes and density can be correlated to their respective per-pixel *r* values and conversely pixels with a given anisotropy can be assigned with a sub-diffraction pixel nano-distribution **(Figure S13).** While we cannot completely discount the possibility of processing artifacts in the registration of the super-resolution STORM with the homoFRET datasets, the fact that the individual point-spread functions do span several adjacent pixels suggests that this effect is likely minimal **(Figure 7 a-e)**.

**Figure 7.**
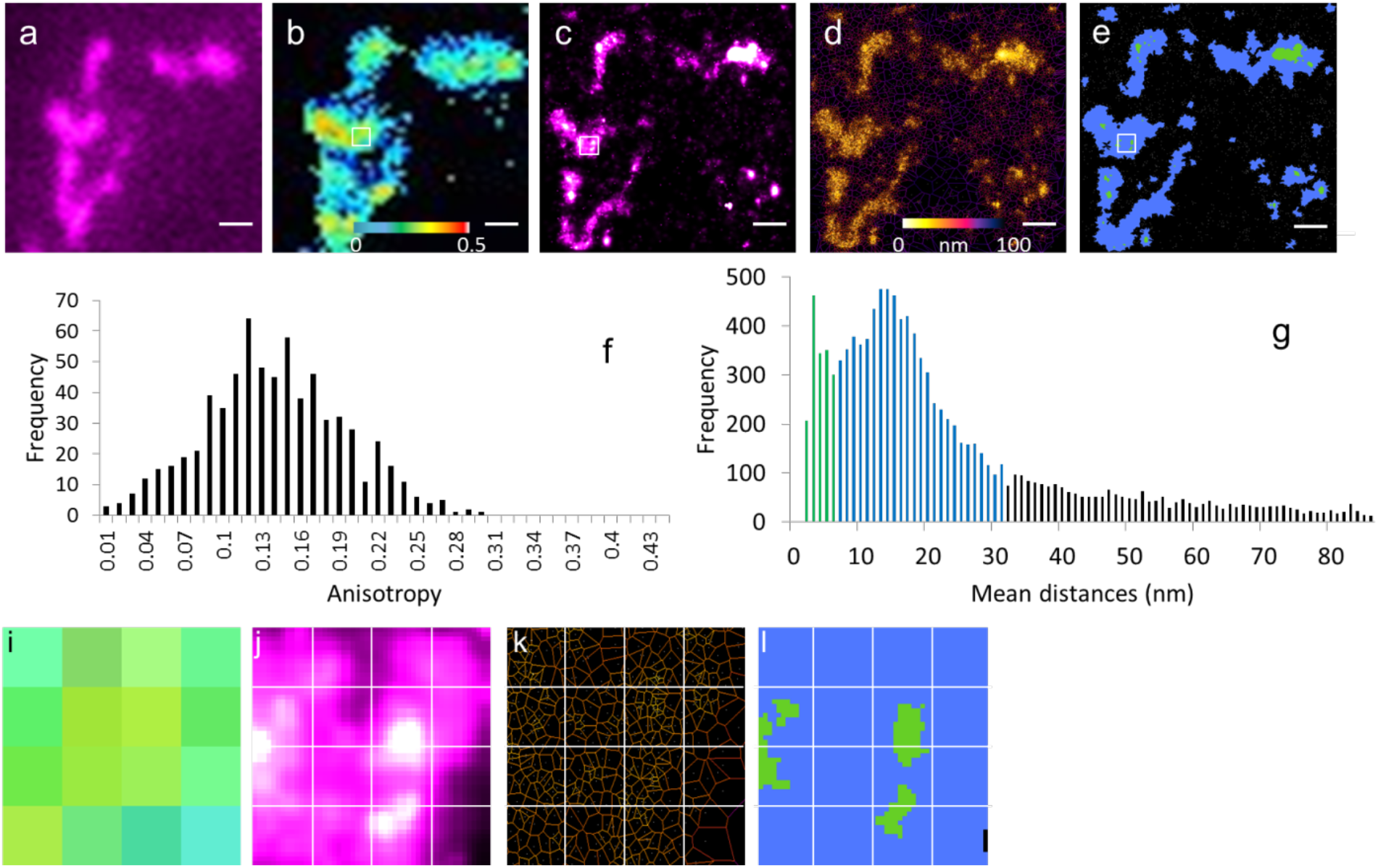
Correlating STORM/homoFRET enables to uncover the preferential spatial distribution of CEACAM1 monomers and oligomers. **(a)** Diffraction-limited image of CEACAM1 labeled with anti-eYFP AF-647 conjugated nanobody. **(b)** HomoFRET anisotropy map of CEACAM1-eYFP. **(c)** Super-resolved image of CEACAM1 labeled with AF-647-conjugated anti-eYFP nanobodies. **(d)** Voronoi tessellation segmentation. **(e)** Segmentation into micro- and nano-clustered areas. **(f)** Histogram of anisotropy values for CEACAM1-eYFP cluster in panel **b**. **(g)** Mean distances distribution derived from Voronoi tessellation segmentation [green - nanoclusters; blue - microclusters; black - non-clustered diffuse regions]. **Panels i-l**: 4 x 4 pixels with a pixel size of 127 nm scaled up to reveal detail within regions denoted by white boxes in panels **b, c and e**, respectively: **(i)** Anisotropy map, **(j)** Super-resolved CEACAM1-AF647 **(k)** Voronoi tessellation segmentation; **(l)** Segmentation into micro- and nano-clustered areas.

What becomes clear from this analysis is that pixels with equal anisotropy values display a wide range of possible nanoscale distributions **(Figure 7 i-k).** It is notable that low anisotropy pixels tended to be devoid of nanoclustered regions, instead appearing to be comprised of micro-clustered and diffuse CEACAM1. These findings suggest that CEACAM1 organizes preferentially into three characteristic states: a diffuse distribution of oligomers, microclusters of oligomers and monomers, and nanoclusters of monomers (**Figure S14**). We note that similar cluster classifications were reported by others studying T-cells using SMLM [39]. In addition to this approach, we have also correlated the anisotropy map to its corresponding nearest neighbour map. Figure S15 shows that the higher anisotropy pixels at the core of the cluster have, on average, localizations with a lower distance to their nearest neighbour. This is consistent with our observation of a negative correlation between self-association and protein density.

## Discussion

The present work has demonstrated that combining STORM with homoFRET imaging enables direct determination of the spatial and oligomeric state distributions of surface membrane proteins. In the case of CEACAM1, we have shown that this glycoprotein can exist in three distinct clustered states, each with its own unique self-association characteristics: diffuse regions consisting of oligomers present on the flatter cell surface, microclusters of oligomers and monomers, and nanoclusters of monomers that tend to be within cell protrusions. These arrangements are not resolvable using conventional diffraction-limited approaches. In the present work, cell size, morphology, or cell cycle were not considered in the analysis, extensions of the approach described here to consider these additional parameters is under consideration.,

While our studies focused on the full-length variant of CEACAM1, the relative ease of implementation both in terms of microscope setup **(Figure 8)** and labeling strategy makes this approach applicable to virtually all membrane proteins. This method can be generalized to other fluorescent proteins. The availability of other anti-xFP-specific nanobodies conjugated to other photoswitchable organic fluorophores spanning the visible spectrum, as well as other fluorescent xFP-mutants, opens the door for multicolour STORM/homoFRET experiments, simultaneously monitoring the arrangement of multiple cellular proteins. We have also highlighted the importance of selecting a suitable cluster analysis strategy in order to categorize single molecule localizations into suitable groups. In our hands, Voronoi tessellation proved to be the most relevant approach as it allows automated subdivision of points into polygonal regions without user bias. Nonetheless, although these determinations were performed without considering localization uncertainty, the observed differences in density are still representative of differences in labeling density that reflect CEACAM1 density. The resulting mean distance or density distributions revealed a trimodal distribution for CEACAM1 in a subset of the analyzed cells, showing three discrete populations that were indistinguishable under diffraction-limited fluorescence microscopy. Having established a system capable of revealing the subtle, nanoscale 2D arrangement of CEACAM1 at the membrane, future studies can now aim to understand how different CEACAM1 isoforms and mutants compare, providing new insights regarding their respective roles and the evolutionary need for such a large array of glycoproteins.

**Figure 8.**
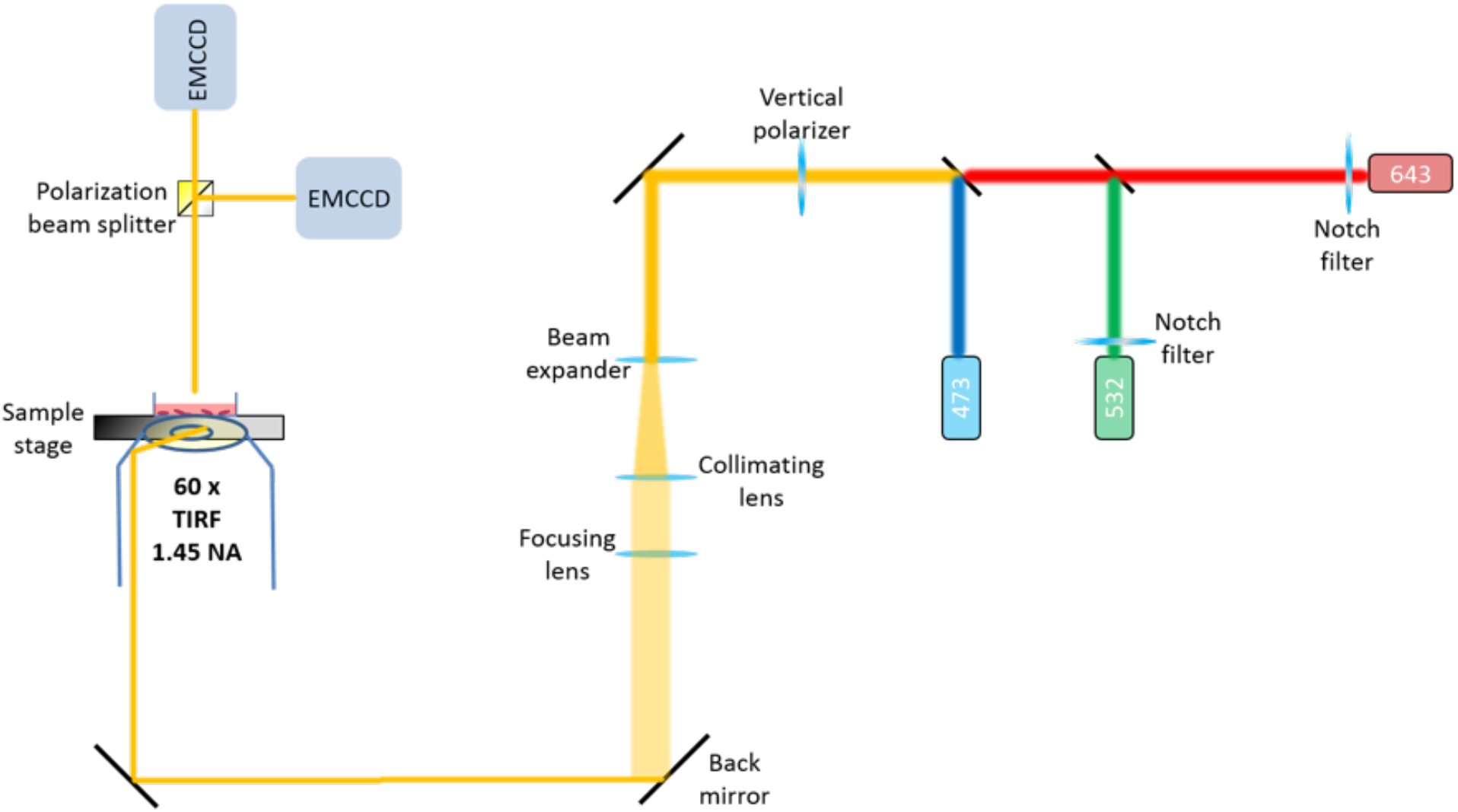
Schematic of the microscope used to conduct STORM/homoFRET experiments.

## Methods

### Cell culture and transfection

HeLa cells were cultured in 75 cm^2^ Falcon culture flasks with supplemented RPMI Medium 1640 (1X) + L-Glutamine (Gibco^®^ by LifeTechnologies™) containing 10% fetal bovine serum (FBS, Gibco^®^ by Life Technologies^™^), 1% glutamax and 0.05% penicillin/streptomycin. Cells were incubated at 37°C with 5% CO_2_. Upon reaching 70-80% confluence, cells were detached using 1x trypsin:PBS for 5 min, and were seeded on Nunc^™^ Lab-Tek^™^ II 8 well chamber slides (Thermo Scientific^™^) with RPMI containing FBS, glutamax, penicillin/streptomycin. Cells were left to adhere and divide for 24-36 h prior to transfection. Transfection with the CEACAM1-eYFP-encoding was conducted by resuspending 10 μL of the plasmid in 87 μL of RPMI-free serum for 5 min, followed by the addition of 3 μL polyethylenimine (PEI) and incubation at room temperature (22°C). 25 μL of the total 100 μL transfection solution is added to each imaging chamber. Transfected cells are incubated for 16-18 h, upon which media is switched to transfection reagent-free media.

### Microscope setup

TIRF microscopy was performed on a home-built TIRF microscopy system integrated with an Olympus FluoView 500 confocal microscope using an IX-70 base (Olympus Canada, Markham, Ontario) using a high numerical aperture 60X oil-immersion objective (NA=1.45, Olympus Japan, Tokyo Japan). A thin layer of index-matching oil (n= 1.518) was used to optically couple the objective to the glass surface of the Nunc^™^ Lab-Tek^™^ II 8 well chamber slides (Thermo Scientific^™^). Excitation of eYFP was achieved using an analog modulated 473 nm diode laser (DHOM-L-150 mW, Suzhou Daheng Optics & Fine Mechanics Co., Ltd., China). Excitation of AF647 was achieved using an analog modulated 643 nm laser (Power Technology, Model LDCU5/A109, Arkansas, USA) with a maximum measured power of 90 mW at the source. A clean-up notch filter (ZET642/20x, Chroma, Bellow Falls, VT) was used to spectrally clean the excitation. A polarization beam splitter was used to separate between polarized and depolarized emission (PBS251, Thorlabs, Newton, NJ) **(Figure 8).** Fluorescent images are captured using two water-cooled eXcelon-equipped Evolve 512 EMCCD camera (Photometrics, Arizona, USA) using μ-Manager (version 1.4.19). Under these conditions, we measure an effective pixel size of 127 nm. Imaging for a single fixed cell was performed over a 30 minute period and 5-10 cells were imaged.

### TIRF homoFRET

TIRF-based homoFRET images are collected as pairs of images where F_ll_ is the polarized fluorescence intensity collected through a polarizer oriented parallel to the excitation and F_⊥_ representing the depolarized fluorescence intensity collected through a polarizer oriented perpendicular to the excitation source. All images were collected using 500 ms exposure period, gain of 1, and an electron multiplier gain of 100. To correct for detector sensitivity for both F_ll_ and F_⊥_ that arise from differences in their transmissivities through the various optical components of the microscope, an experimental G factor is determined. Once the G factor is determined, anisotropy (*r*) can be determined through the following equation:

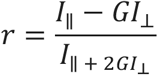

The G factor value of 0.93 was determined by calculating the intensity ratio of I_ll_ / I_⊥_ for a 100 nM fluorescein standard solution. G factor measurements are performed whenever an optical component is modified or moved into the optical train of the microscope.

### Preparation of labeled nanobodies

Nanobodies were purchased from Chromotek as GFP-Trap. GFP-Trap nanobodies were conjugated to AF647 using conventional conjugation protocols. For labeling, 50 μl of the 1 mg/ml protein solution was dialyzed into 0.2 M NaHCO_3_, pH 8.2 using a mini-dialysis unit (Pierce, molecular weight cutoff = 3,500 Da). Succinimidyl-ester of AF647 was dissolved at 10 mg/ml in DMSO, which was added in excess to the nanobodies. The solution was incubated for 1 h at room temperature. Excess dye was removed via buffer exchange into PBS using a desalting column. Conjugated nanobodies were stored at 4°C.

### Colocalization of eYFP to nanobody-AF647

Colocalization analysis was conducted using the JACoP (Just Another Colocalization Plugin) ImageJ plugin. Analysis was conducted on 6 cells from 3 independent experiments. Costes automatic thresholding was used prior to calculating the Pearson coefficient between eYFP and AF647 images[45]. Arithmetic mean and standard error of the mean were calculated and shows the high degree of colocalization between eYFP and AF647.

### STORM imaging

In order to initiate stochastic photoswitching for STORM, photoswitching buffer is added prior to imaging. The buffer consists of 50 mM cysteamine (2-mercaptoethylamine, Sigma-Aldrich), 40 μg/ml catalase (from bovine liver, aqueous solution, Sigma-Aldrich), 0.5 mg/ml glucose oxidase (*Aspergillus niger*, Sigma-Aldrich), 50% w/v glucose (D-glucose, Sigma-Aldrich) diluted in pH 7.4 PBS buffer. This buffer provides conditions that yield a high photon count for AF647, and modulates the photophysical properties of AF647 by scavenging oxygen and creating a reducing environment. A 643 nm laser is set to a power of 20 mW, as measured after the objective, which is used to drive AF647 to an off-state prior to having subsets of fluorophores coming back on in a stochastic manner over the acquisition period. To reconstruct a super-resolved image, 10 000 to 15 000 images were acquired, each with an exposure time of 30 ms.

### STORM processing

Images are processed using ThunderSTORM (version 1.3) and the maximum likelihood localization algorithm (MLE) localization parameter. Following localization and reconstruction, the coordinates of single emitters are filtered based on their localization precision (uncertainty value) and photon count in order to discard electronic noise. Specifically, uncertainty histograms are generated for each cell and experiment. We found that thermal and electronic noise resulted in uncertainities of [0-6] nm while sample noise affected most localizations with a localization precision above 40 nm. Localizations with a neighbour within ~ 1 nm are merged into a single localization in order to correct for self-clustering artifacts arising from multiple blinks of the same fluorophore or dual labels on a nanobody. Despite the care taken in accurately pre-processing and post-processing blinks, and due to the stochastic nature of photoswitching, the variation in the number of fluorophores per protein (albeit small) and overall imaging conditions, the super-resolved localization coordinates do not offer an absolute measurement (i.e: the ability to count the number of proteins in a region) of protein count but a rather a statistically robust, relative quantitative representation of protein distribution and clustering.\

### STORM cluster analysis

Several clustering algorithms have been used by other groups on sets of coordinates obtained from SMLM techniques with a preference for Ripley’s K function, pair-correlation analysis and density based spatial clustering analysis with noise (DBSCAN) [39, 40]. While useful, these methods present some limitations as they require user-assigned analysis parameters, which will skew the resulting clustering analysis [40–42]. Here we use Voronoi tessellation, which is a geometric method that subdivides space into polygonal areas [43]. The region covered by all the polygons forms a mosaic of tiles of varying sizes that define what is called as a Voronoi diagram. In this approach, each polygon is centered around one localization, hence describing single molecules neighborhoods with their respective properties. Since a single STORM reconstructed image would contain 10^5^ to 10^6^ localizations depending on the sample, SR-Tesseler uses a sweep line algorithm to generate a Voronoi diagram in a matter of seconds (33 seconds for a full 512 x 512 pixels field of view with approximately 400,000 localizations). We use the SR-Tesseler software for this purpose. Note that localization uncertainty is not weighted into the cluster analysis.

### Nearest neighbour analysis to characterize the median distance between nearest neighbour localizations

Each localization is assigned a nearest neighbour defined as the closest localization in the super-resolved dataset. Localization uncertainties are not weighted in the distance calculation (code available in supplementary information).

## Acknowledgments

We thank Dr. Uros Kuzmanov for his generous gift of single domain antibodies and Maximiliano Giuliani for writing the nearest neighbour analysis code. This work was funded by Canadian Institutes of Health Research Operating Grant: MOP-123320 and the Natural Sciences and Engineering Research Council of Canada Discovery Grant: RGPIN-2015-043. We thank John Oreopoulos, Jocelyn Lo and Kelsey Downie for previous work on this project. We thank Professor Jonathan Rocheleau, University of Toronto for the gift of the monomeric and dimeric Venus constructs. We also thank Nelly Leung for her assistance in cell culture.

## Supplementary information

### Method development

**Figure S1.**
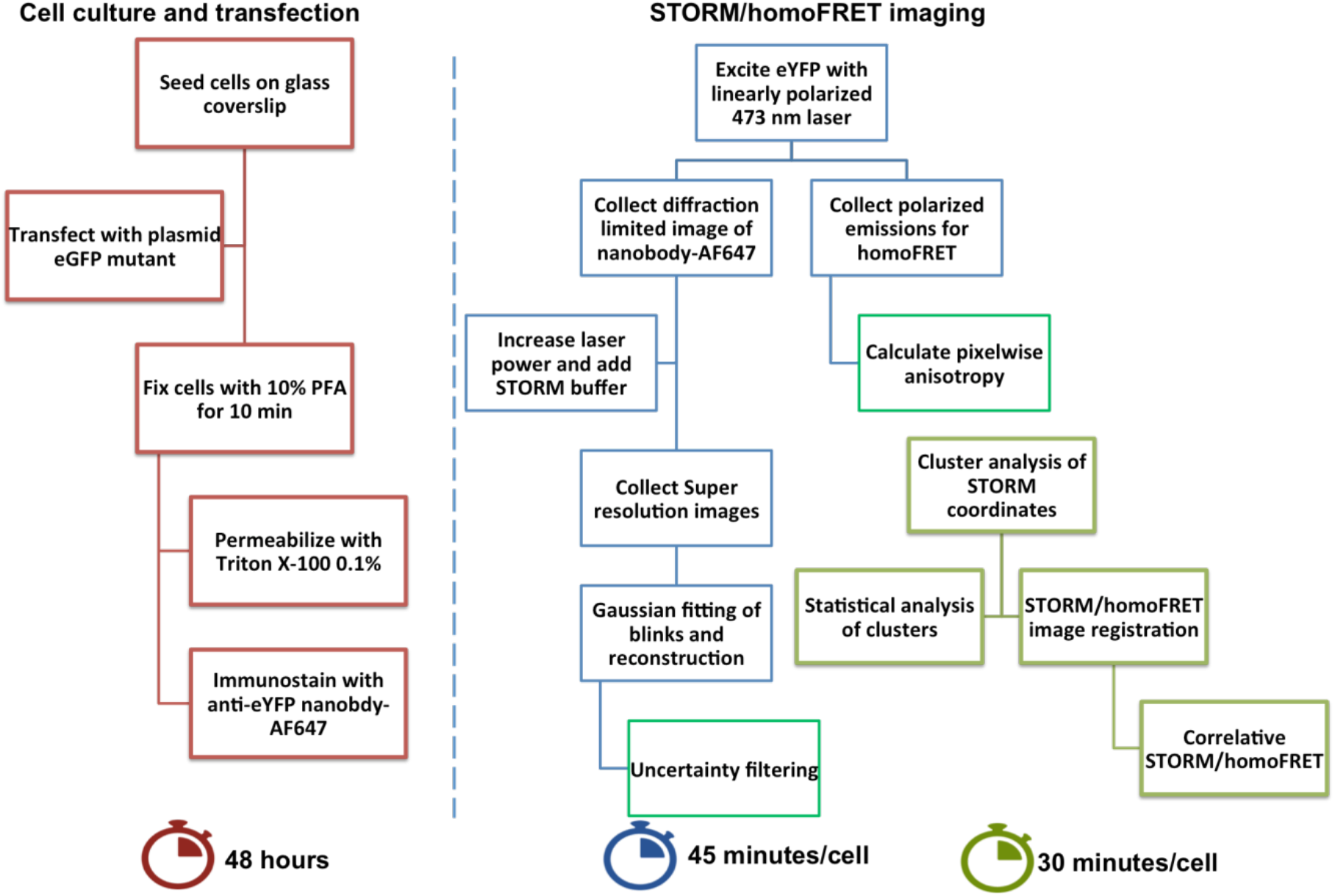
Workflow required to perform STORM/homoFRET measurements in cells transiently transfected with an eYFP fluorescently labeled protein, and immunolabeled using an anti-GFP single domain antibody.

**Figure S2.**
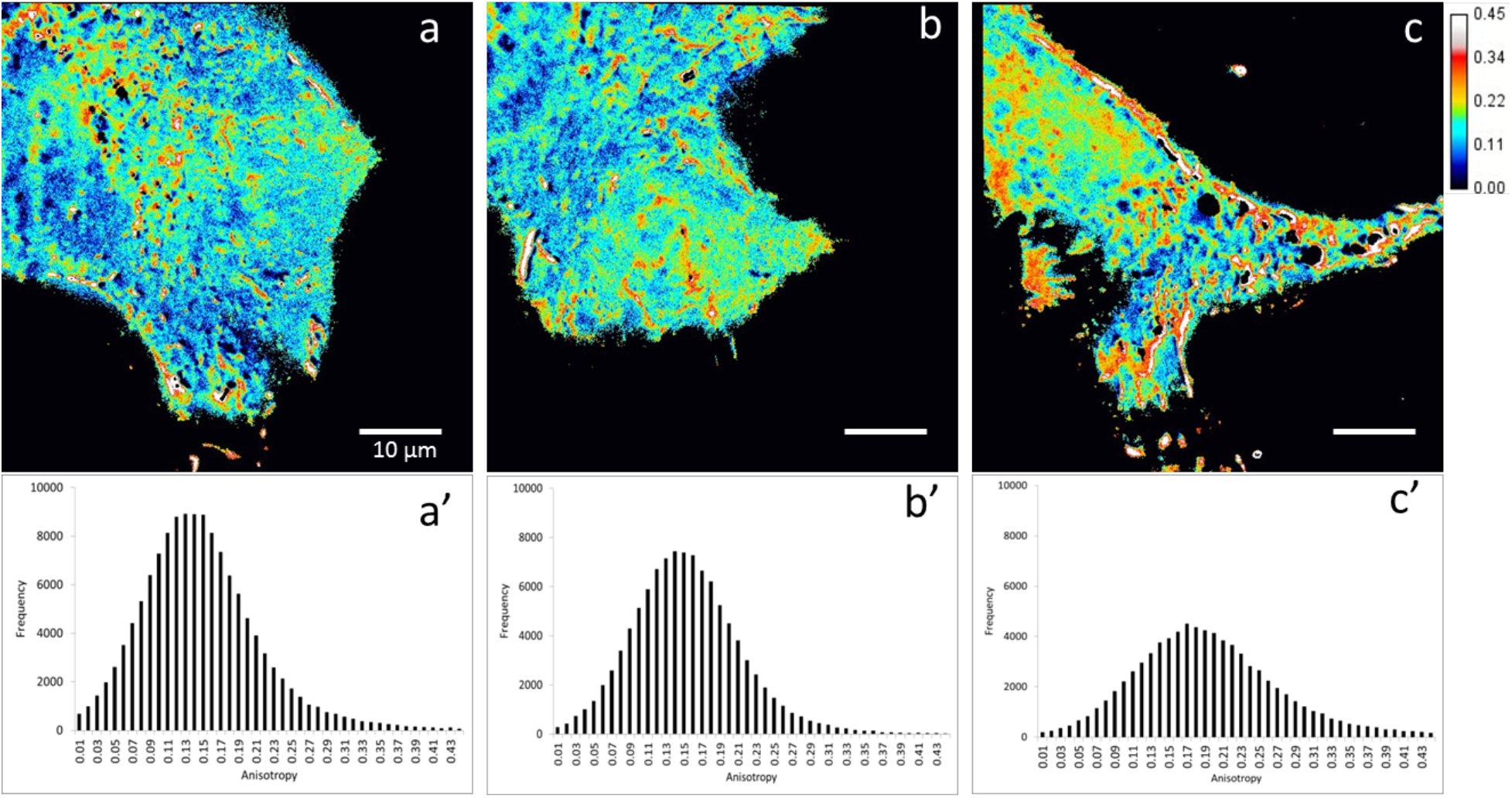
Different cells exhibit slight variation in their distribution of monomers and oligomers. However, the mixture of oligomeric forms remains a common attribute of all CEACAM1-eYFP transfected cells. (**a, b, c)** Anisotropy maps from homoFRET measurements displaying a heterogeneous distribution of CEACAM1 monomers and oligomers. **(a’, b’, c’)** Corresponding anisotropy distributions for each of the associated images.

**Figure S3.**
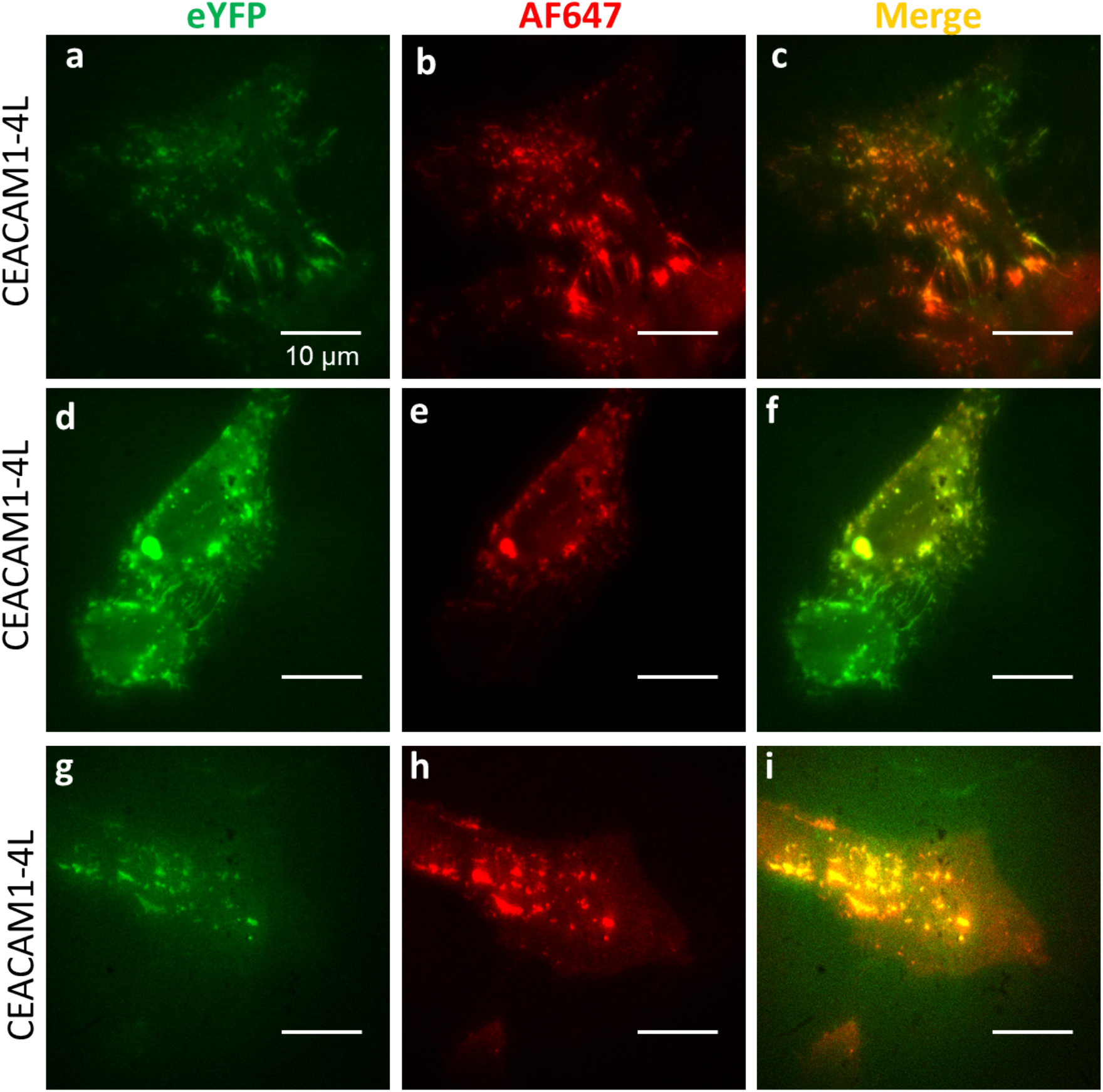
Representative images of the colocalization analysis showing the specificity and accuracy of the single-domain antibody labeling. **(a, d, g)** Transiently transfected HeLa cells with CEACAM1-eYFP, **(b, e, h)** Single domain antibody-AF647 labeled cells, **(c, f,i)** Merged eYFP and AF647 channels display the extent of colocalization.

### Validation of homoFRET

In order to validate our TIRF-homoFRET measurements, we compared the steady-state anisotropy values for monomeric and tandem-dimeric Venus fluorescent proteins **(Figure S4**. HeLa cells were transiently transfected with either Venus monomer or Venus tandem-dimer. Cell media was exchanged 18 hours after transfection and the cells were imaged using our TIRF-homoFRET setup.

**Figure S4.**
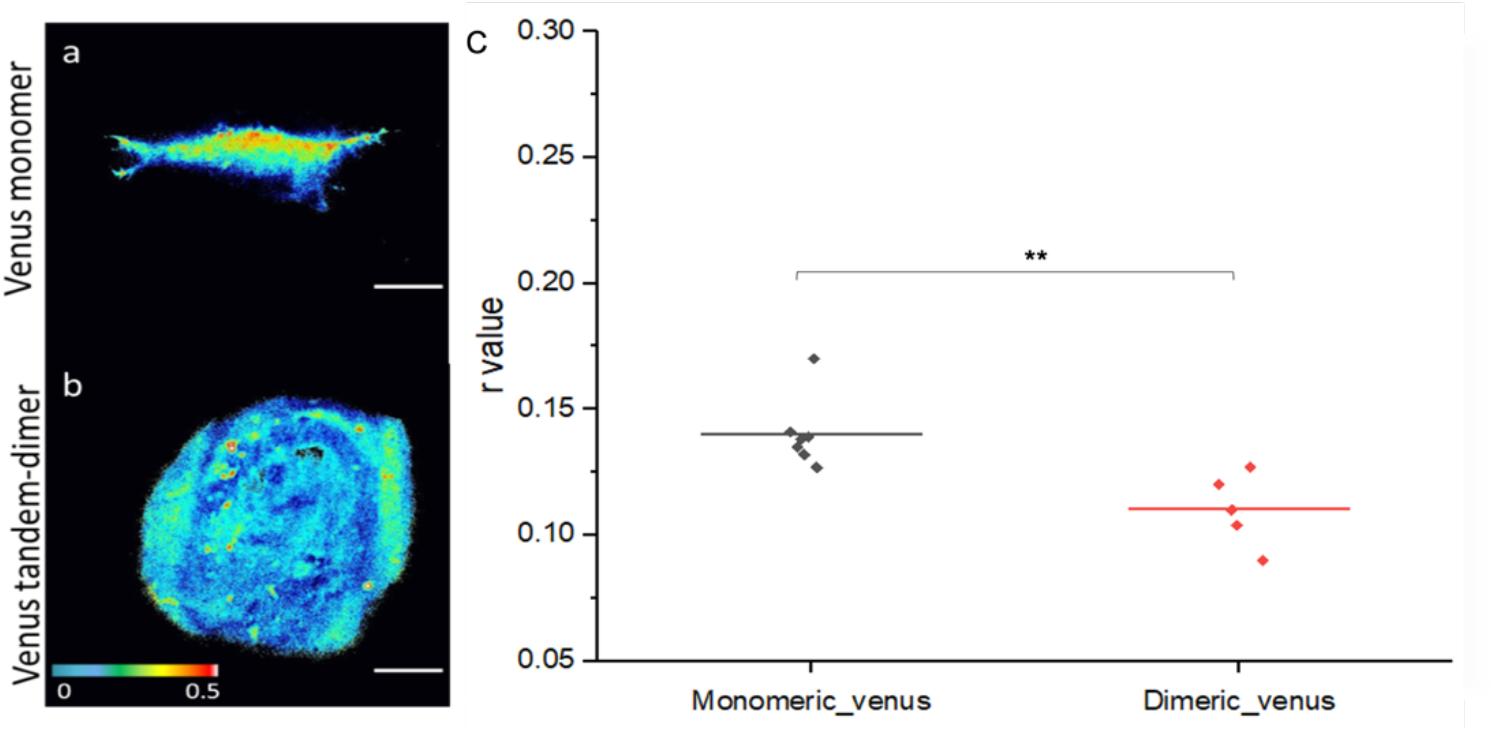
HomoFRET anisotropy control experiment. Representative cells showing the pixelwise anisotropy map for **(a)** monomeric-Venus and **(b)** tandem-dimer Venus transiently transfected in HeLa cells, **(c)** Average anisotropy plots generated from 7 and 5 cells respectively of the monomeric and tandem-dimeric Venus from 2 replicate experiments. Dots represent individual cells. Assuming a normal distribution, using a two-sample t-Test, we find that p < 0.005.

**Figure S5.**
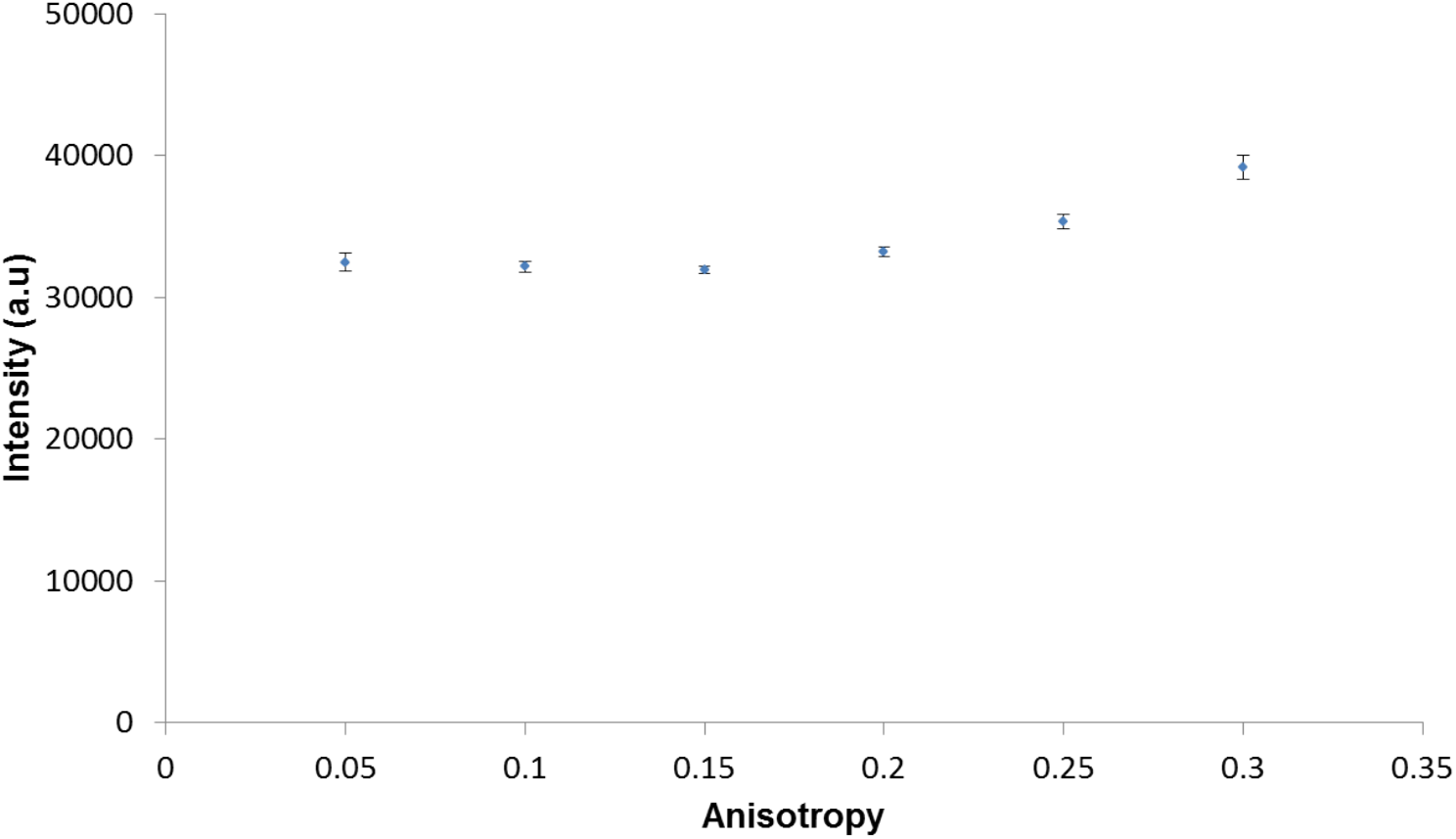
Anisotropy vs Intensity. The relationship between pixel-wise anisotropy and intensity shows that reported anisotropy values are not skewed by intensity but instead represent the ratiometric measurement of anisotropy. Error bars represent standard error of the mean from 6 cells from 2 biological replicates.

**Figure S6.**
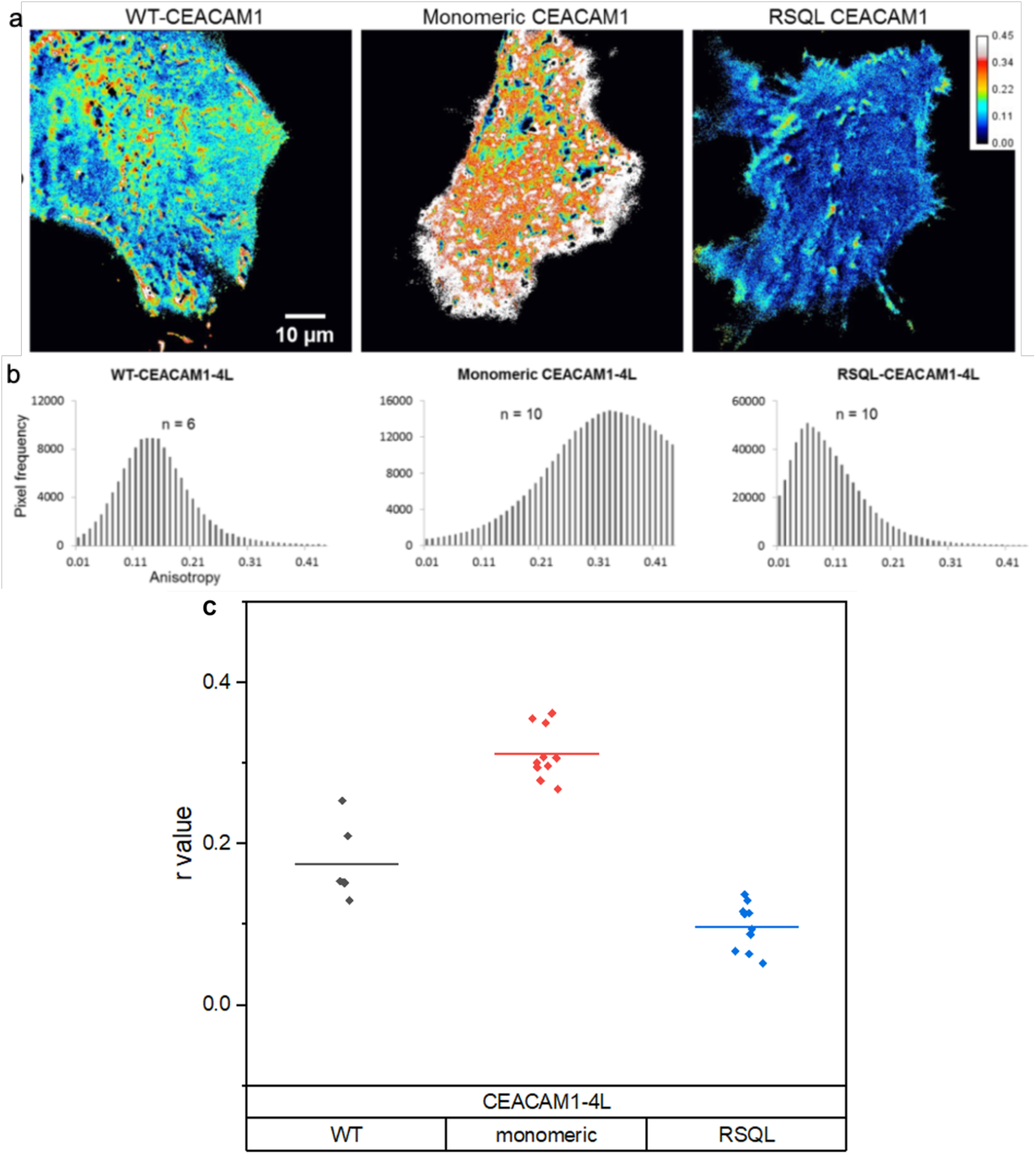
HomoFRET measurements of CEACAM1 WT and mutants shows the validity of the method in capturing expected self-association states. **(a)** Representative images for WT_CEACAM1-4L, monomeric CEACAM1-4L and RSQL-CEACAM1-4L, which is unable to form trans-homophilic interactions, **(b)** Cumulative histograms of intracellular anisotropy distributions for each of the CEACAM1 variants. Each n represents an individual cell and the histograms represent the sum of the individual histograms from each cell.

**Figure S7.**
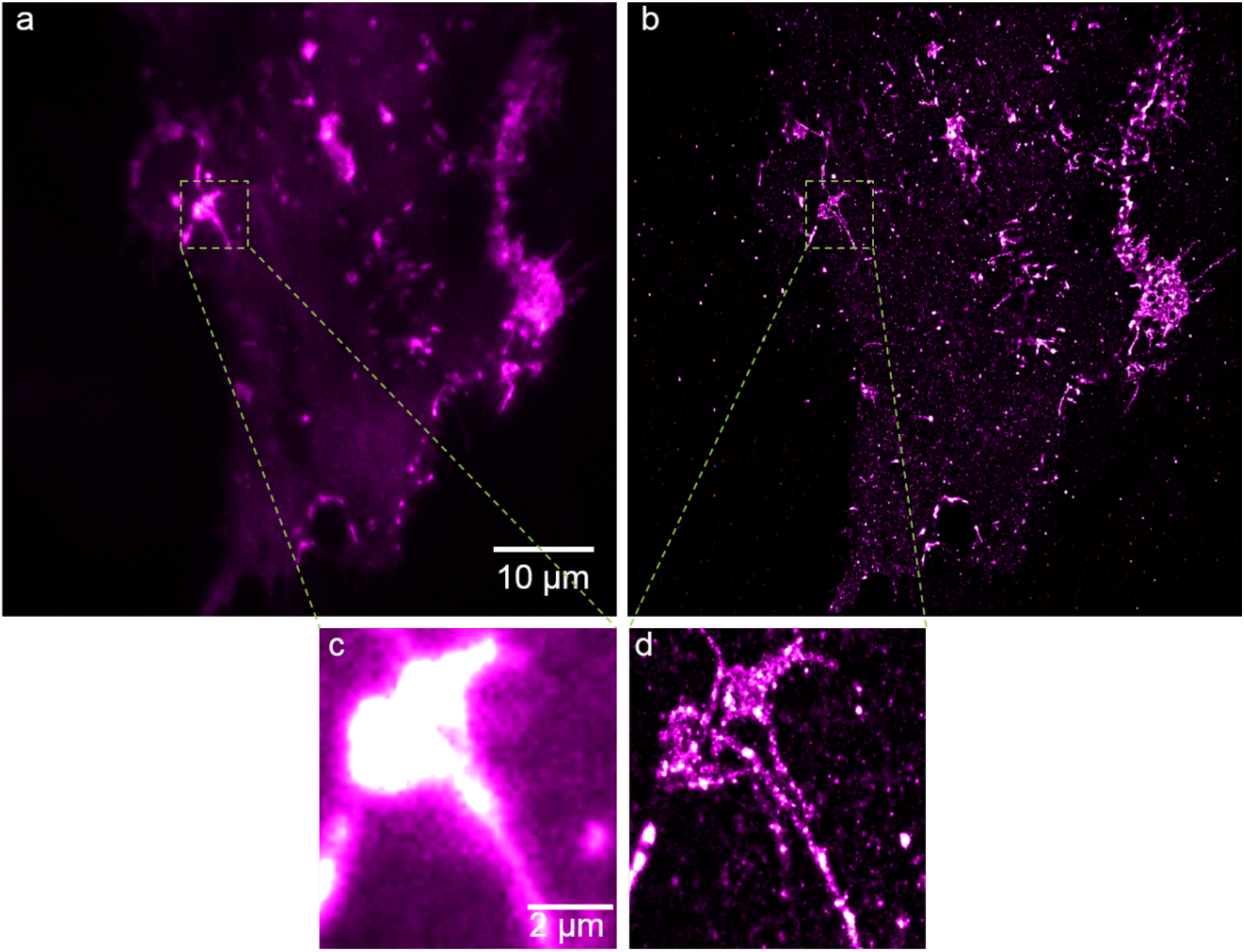
Diffraction-limited and super-resolved CEACAM1-4L **(a)** Diffraction-limited image of CEACAM1-4L transiently transfected in HeLa cells, **(b)** Superresolved image of CEACAM1-4L transiently transfected in HeLa cells, **(c)** Diffraction-limited CEACAM1-4L cluster appears as a homogeneous entitity based on intensity, **(d)** Super-resolved image of CEACAM1-4L cluster reveals a heterogenous protein distribution.

**Figure S8.**
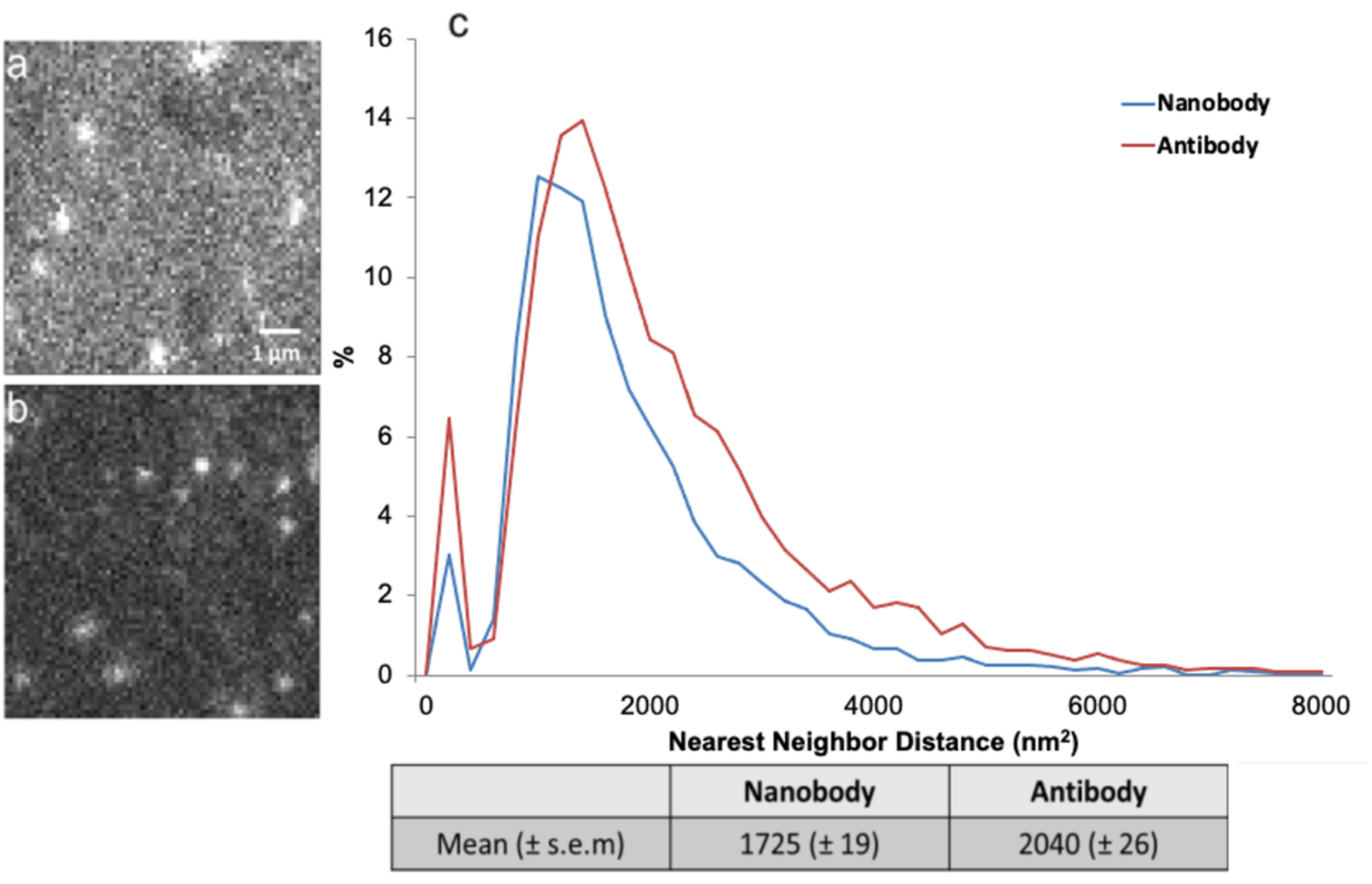
PSF overlap comparison between antibody and nanobody labeling strategies. **(a)** Representative image showing blinks generated in a STORM experiment using antibodies (pixel size 127 nm), **(b)** Representative image showing blinks generated using nanobodies, **(c)** Histogram of nearest neighbour distance from blinks in individual frames. 10 frames per stack of 5000 frames taken at 500 frame intervals for each cell with n= 3 cells from independent experiments for each of the labeling strategies. Note that representative images are shown with adjusted contrast so as to display the lower signal to noise ratio in nanobody-labeled images that arises from the lower number of fluorophores per diffraction-limited region.

Since the AF647-labeled nanobody is bound to eYFP, which is itself bound to a flexible linker attached to the cytoplasmic domain of CEACAM1, we expect that there is a small, yet non-negligible fluctuation of the fluorophore’s position over the acquisition time. If not accounted for, this might lead to the localization of blinks at slightly different positions for the same fluorophore, which would lead to an overestimation of cluster density. Note that the baseline fluctuation arising from the thermal noise of the EMCCD and laser fluctuation was calculated in order to establish the median instantaneous fluctuation of both eYFP and AF647. Here we characterize the median fluctuation of both eYFP and AF647 in order to establish a lower bound for STORM’s localization precision. For this purpose, we captured diffraction-limited images of both eYFP and AF647 with an exposure of 20 ms and a time interval of 30 ms. A Gaussian is fit iteratively to isolated PSFs inside the cell in order to measure the displacement in x and y.

**Figure S9.**
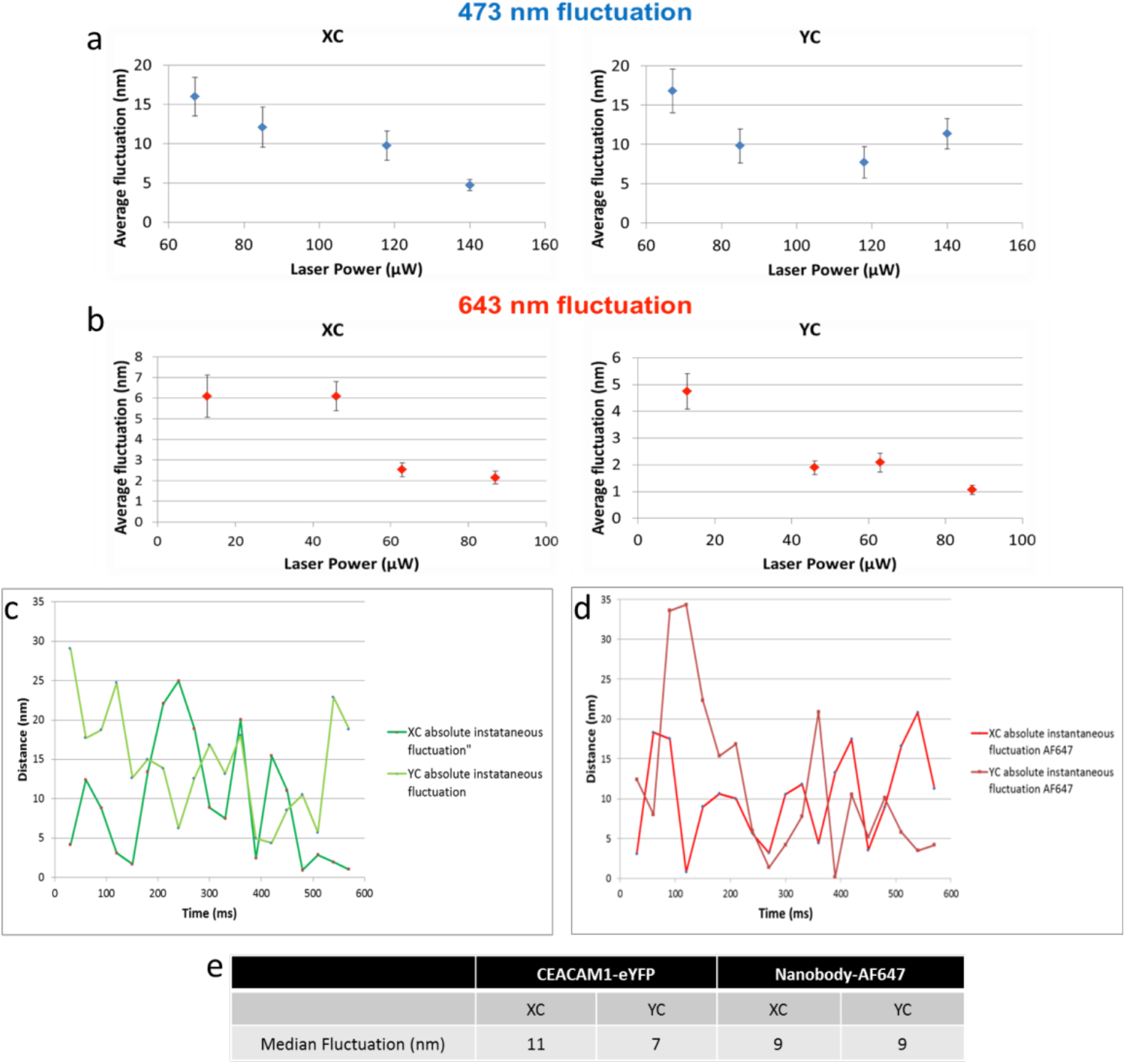
Measure of average **(a)** 473 nm and **(b)** 647 nm fluctuation as a function of laser power. Fluorescent microbeads were imaged in TIRF and the fluctuation was measured by determining the centroid of the bead over time. This experiment shows that intensity fluctuation decreases with higher laser power. **(c)** eYFP absolute instantaneous fluctuation, **(d)** AF647 absolute instantaneous fluctuation, **(e)** Median fluctuation of eYFP and AF647-nanobody conjugates in HeLa cells transiently expressing CEACAM1-4L.

**Figure S10.**
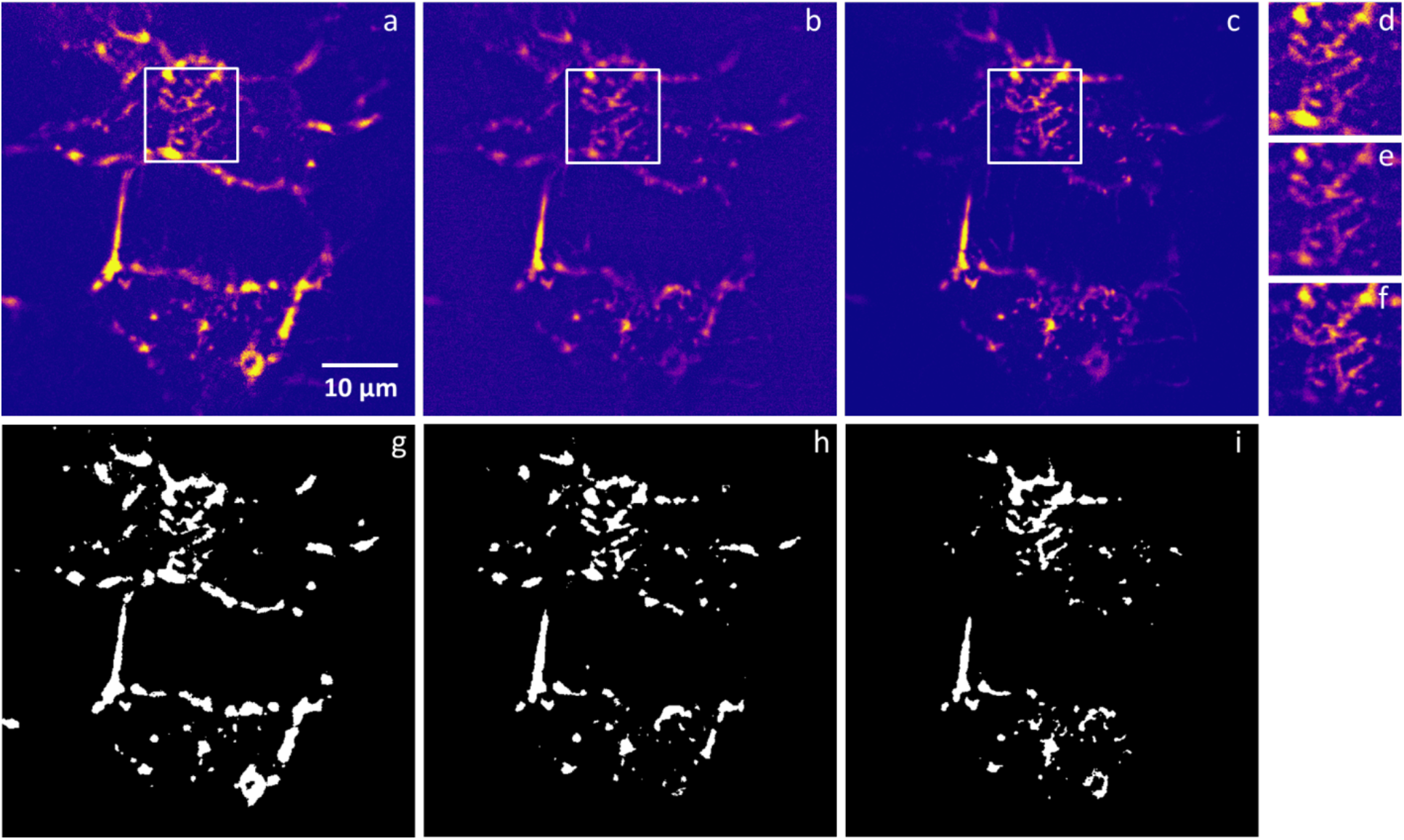
**(a)** HeLa cells transiently expressing CEACAM1-4L fixed with 4% PFA, **(b)** cell permeabilized using 0.1% Triton X-100, **(c)** nanobody labeled cell. Selected ROI for **(d)** fixed, **(e)** permeabilized and **(f)** nanobody-labeled cells. Binary images of **(g)** fixed, **(h)** permeabilized and **(i)** nanobody-labeled cells, respectively. Binary images were obtained by thresholding relative to the fixed-cell image. Certain clusters and regions appear to have disappeared for this reason due to photobleaching.

**Figure S11.**
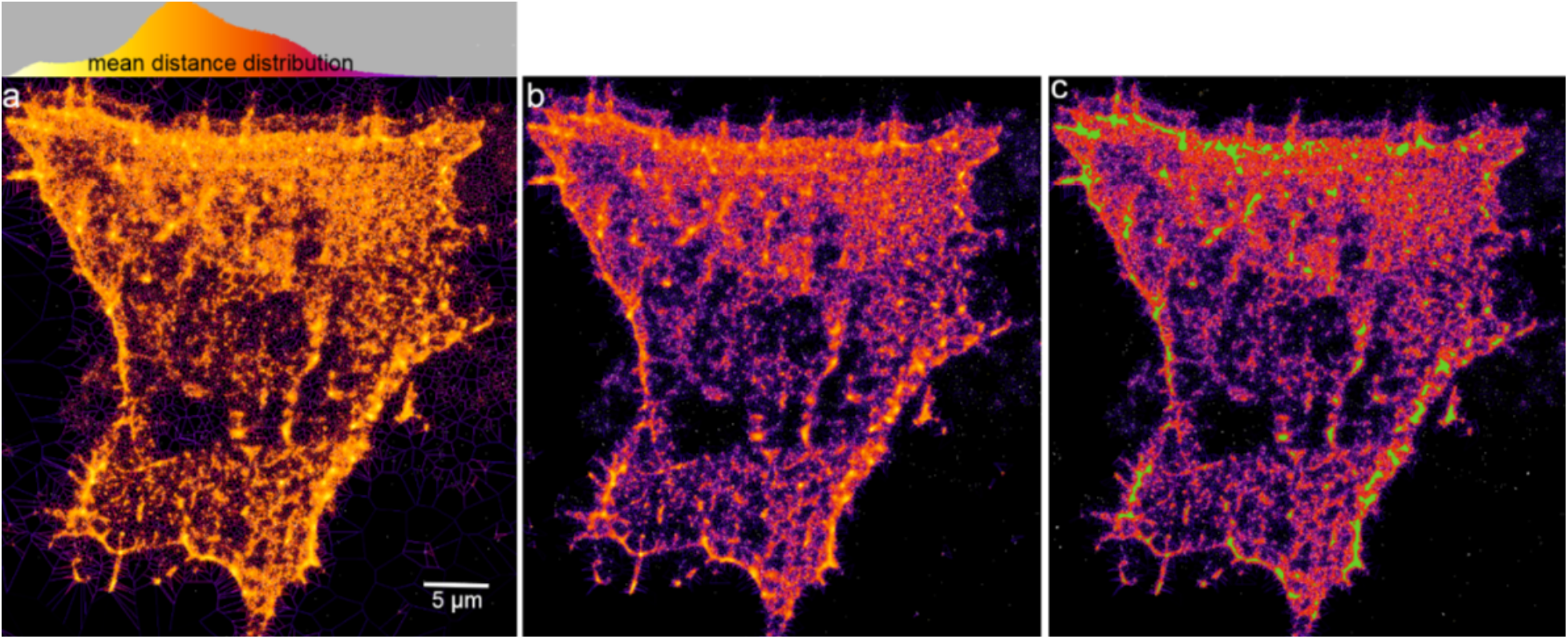
Application of Voronoi tessellation on STORM generated localizations of CEACAM1-4L. **(a)** Voronoi map of all localizations, colour coded for mean distances, (**b)** Voronoi map with background localizations thresholded out, **(c)** Example of clusters (green) that can be segmented using Voronoi tessellation.

**Figure S12.**
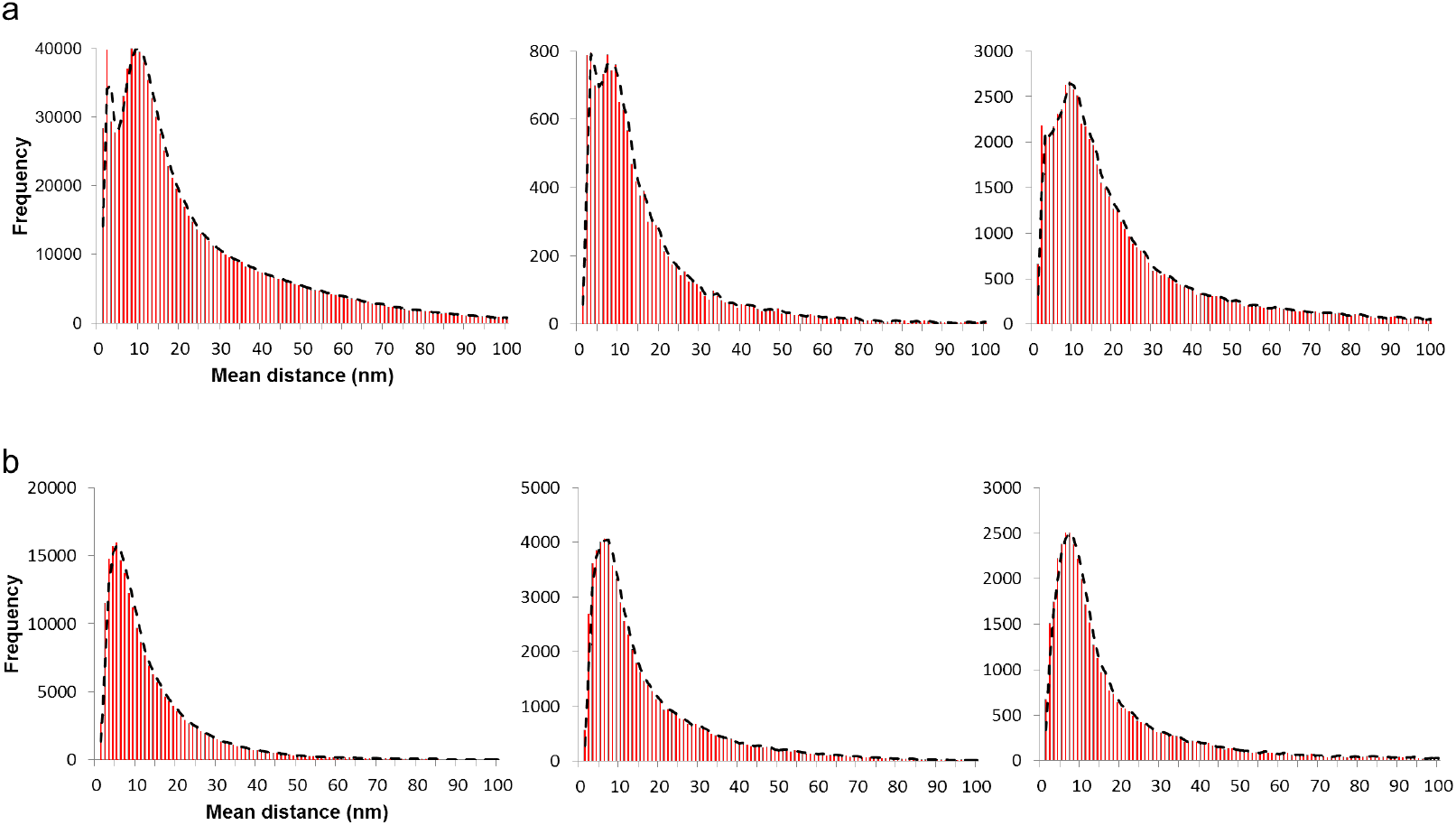
Voronoi tessellation analysis of CEACAM1 highlights that CEACAM1 can arrange its clustering properties depending on the cell’s signaling requirements. **(a)** Three (3) mean distance distributions from 3 cells from 2 replicate experiments showing that CEACAM1 can cluster into as 3 distinct populations, namely: nanoclusters, microclusters and diffuse regions, **(b)** However, in some cells, the nanocluster population is barely resolvable as shown here where only microcluster and diffuse regions can be extracted. Black dotted line represents the moving average (moving average of 2 nm).

**Figure S13.**
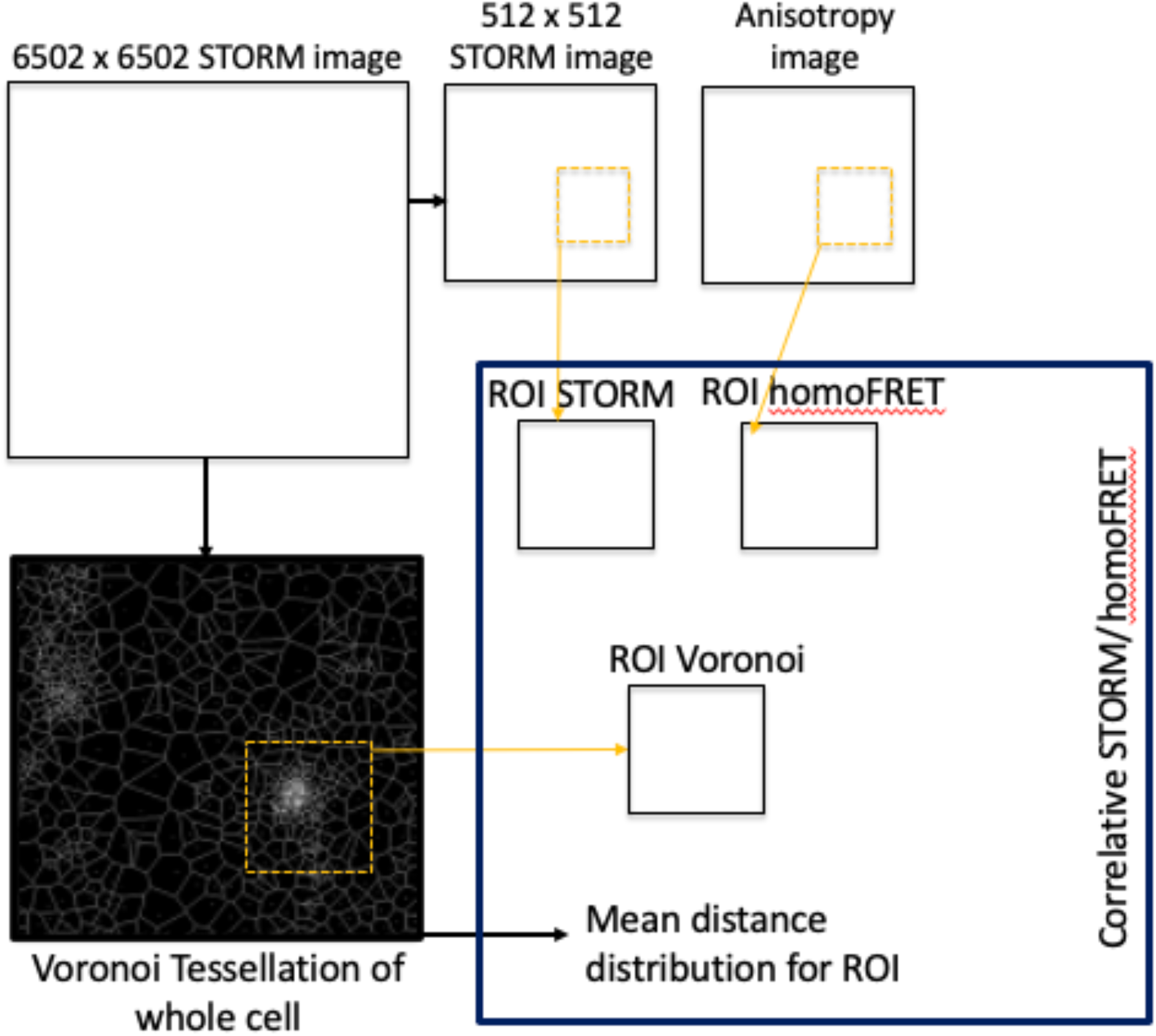
Visual representation of the STORM/homoFRET analysis pipeline. The superresolved image is rescaled without interpolation into a 512 x 512 image while the coordinates are inputted for cluster analysis using Voronoi Tessellation. ROIs are selected for each the STORM image, anisotropy image and Voronoi segmentation. From these ROIs, we obtain the mean distance statistical distribution and the anisotropy distribution.

**Figure S14.**
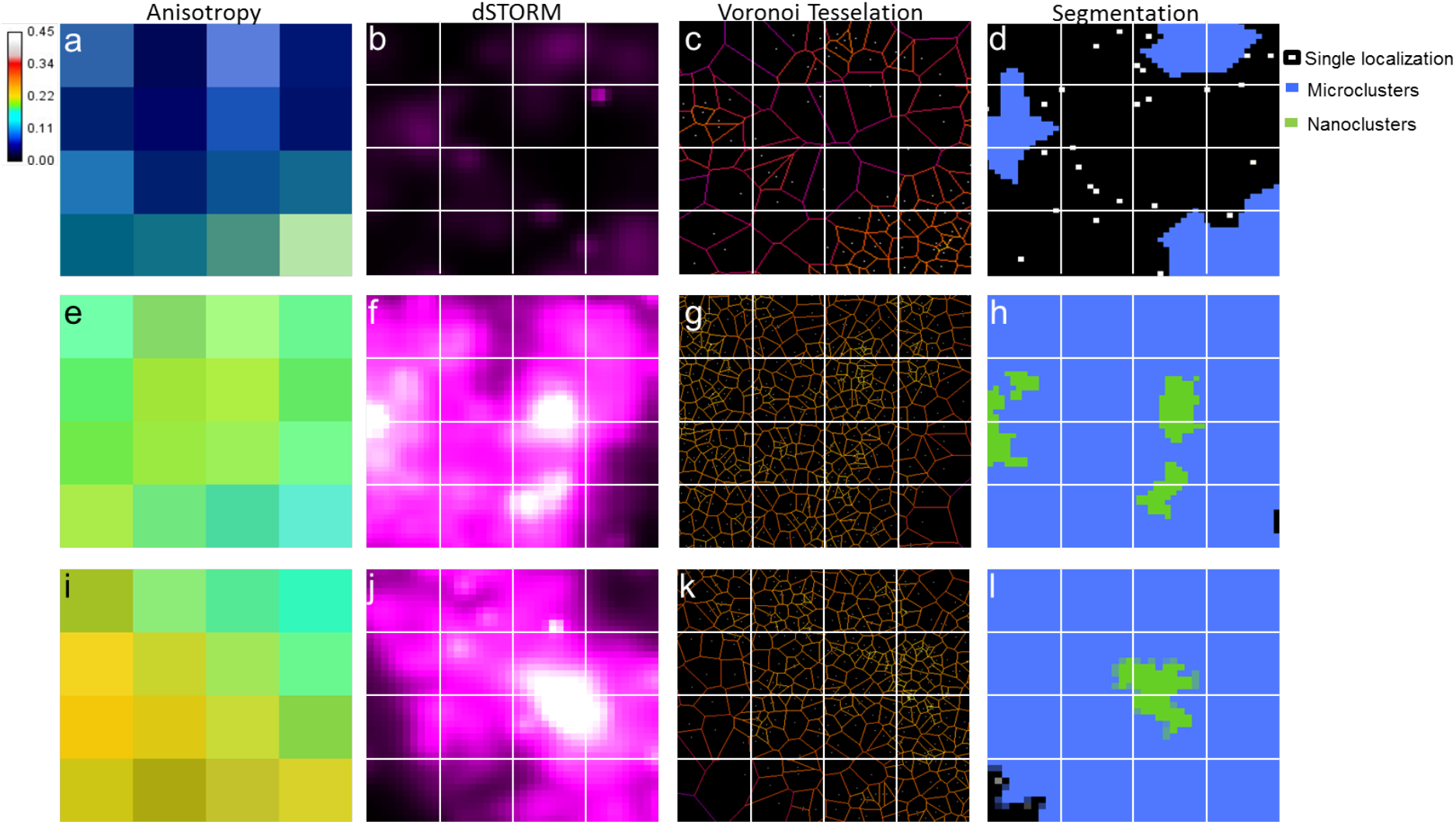
Three distinct ROIs within the same CEACAM1 cluster showing how different anisotropy values correlate with nanoscale spatial distributions obtained via STORM and analyzed through the use of Voronoi tessellation. White lines delimitate diffraction-limited pixel size, with 4 x 4 pixels of 127 nm. **(a)** Anisotropy map CEACAM1-eYFP. **(b)** Super-resolved CEACAM1-AF647. **(c)** Voronoi tessellation clustering. **(d)** Segmentation of the ROI into diffuse and micro-clustered areas. **(e)** Anisotropy map of CEACAM1-eYFP. **(f)** Super-resolved CEACAM1-AF647. **(g)** Voronoi tessellation clustering. **(h)** Segmentation into micro- and nanoclustered areas. **(i)** Anisotropy map of CEACAM1-eYFP. **(j)** Super-resolved CEACAM1-AF647. **(k)** Voronoi tessellation clustering. **(l)** Segmentation into micro- and nano-clustered areas.

**Figure S15.**
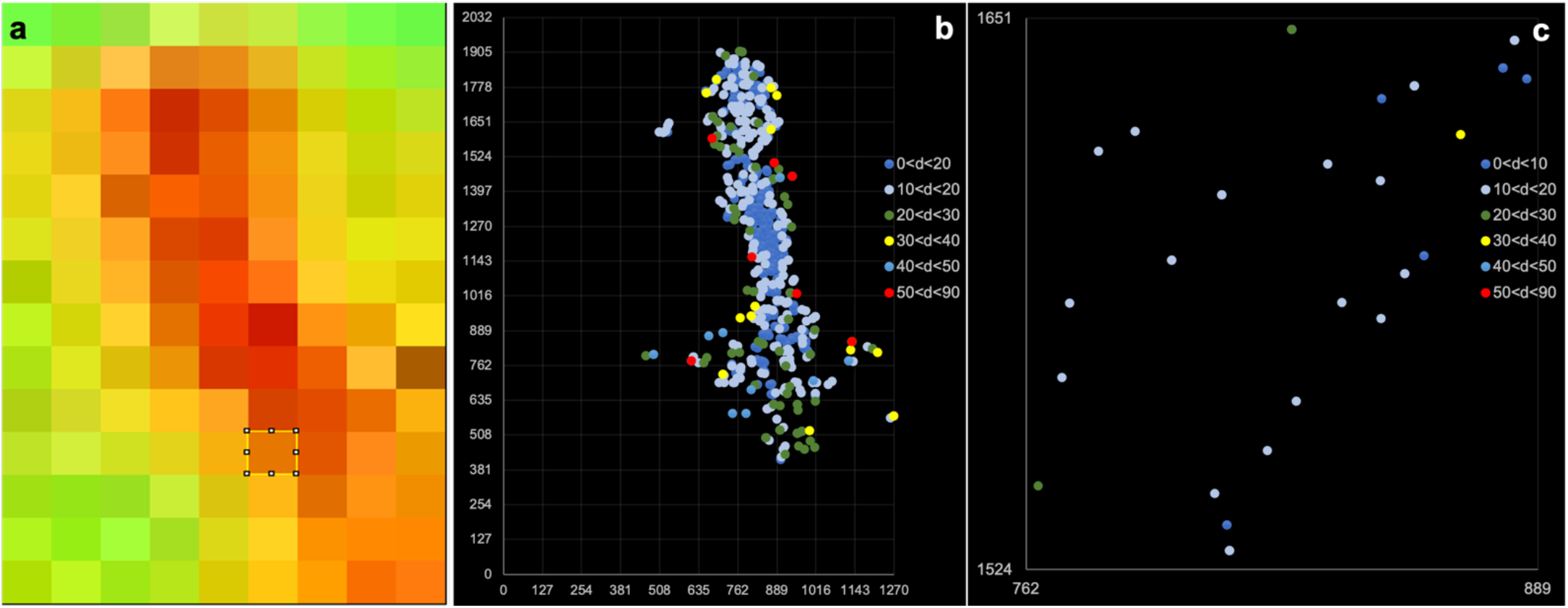
Nearest-neighbour homoFRET/STORM correlation. **(a)** Anisotropy map of a CEACAM1 cluster, **(b)** Color-coded STORM localizations based on NN distances. **(c)** Example ROI (selected box in **(a)** showing individual CEACAM1 localizations (without taking into consideration localization uncertainty).

**Figure S16.**
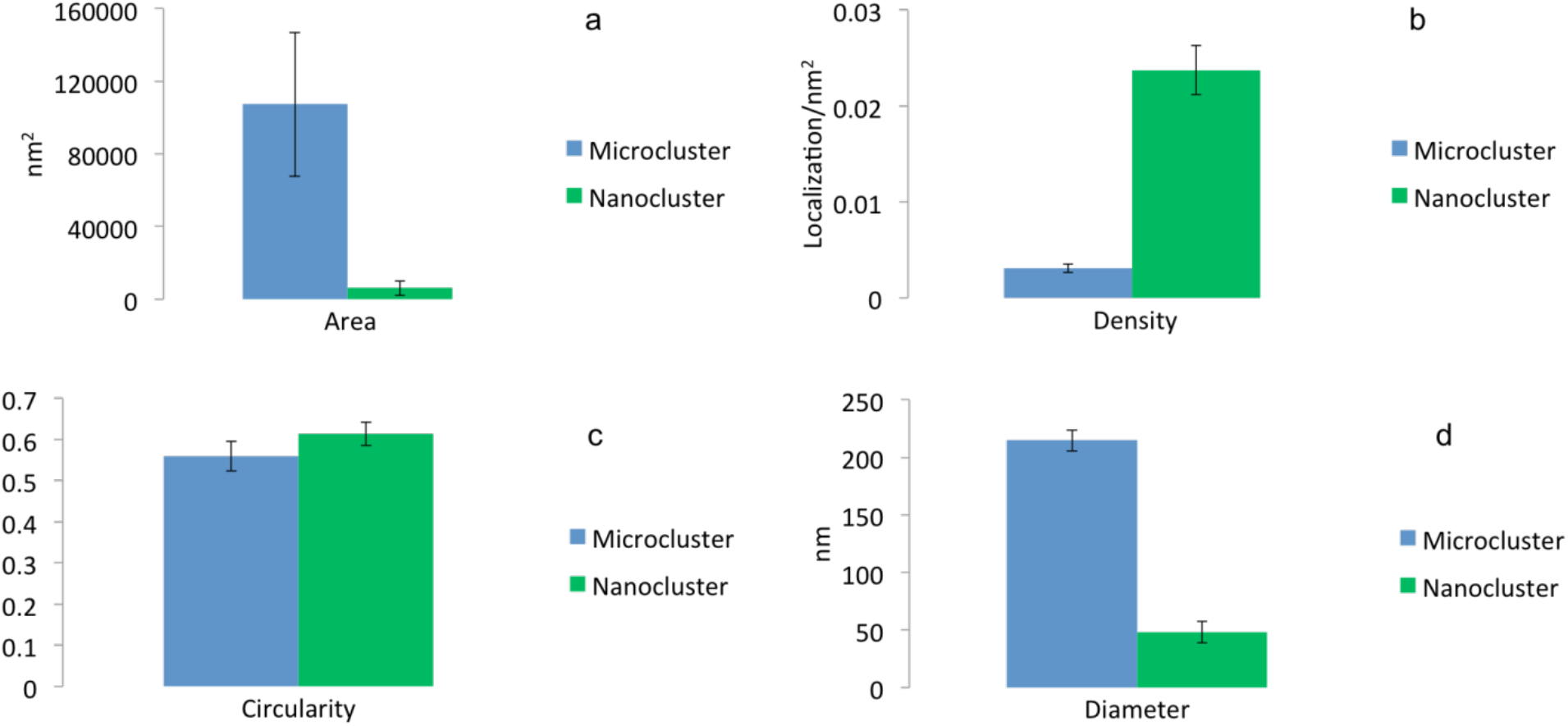
From the various classes of clusters, we can determine their characteristics. Characterization of average cluster area **(a),** density **(b),** circularity **(c)**, and diameter **(d)**, per cluster type, extracted from 8 cells from 3 biological replicates. Error bar represents standard deviation of all clusters per cluster type.

